# The balance between growth and resistance is shifted to the latter by over-accumulation of chloroplast-nucleus located WHIRLY1 in barley

**DOI:** 10.1101/2023.02.22.529264

**Authors:** Monireh Saeid Nia, Susann Frank, Anke Schäfer, Christine Desel, Maria Mulisch, Ulrike Voigt, Daniela Nowara, Yudelsy Antonia Tandron Moya, Wolfgang Bilger, Nicolaus von Wiren, Götz Hensel, Karin Krupinska

**Author notes:** corresponding author: Karin Krupinska. Centre for Plant Genome Engineering (CPGE), Institute of Plant Biochemistry, Heinrich-Heine-University, Düsseldorf, Germany. Institute of Plant Pathology, University of Bonn, Germany (Daniela Nowara).

## Abstract

WHIRLY1 is a chloroplast-nucleus located DNA/RNA-binding protein with functions in development and stress tolerance. By overexpression of *HvWHIRLY1* in barley, lines with a 10-and two lines with a 50-fold accumulation of the protein were obtained. In these lines, the relative abundance of the nuclear form exceeded that of the chloroplast form indicating that over-accumulating WHIRLY1 exceeded the amount that chloroplasts can sequester. Growth of the plants was shown to be compromised in a WHIRLY1 abundance-dependent manner. Over-accumulation of WHIRLY1 in chloroplasts had neither an evident impact on nucleoid morphology nor on the composition of the photosynthetic apparatus. Nevertheless, oeW1 plants were found to be compromised in the efficiency of photosynthesis. The reduction in growth and photosynthesis was shown to be accompanied by a decrease in the levels of cytokinins and an increase in the level of jasmonic acid. Gene expression analyses revealed that already in non-stress conditions the oeW1 plants had enhanced levels of pathogen response (PR) gene expression indicating activation of constitutive defense. During growth in continuous light of high irradiance, *PR1* expression further increased in addition to an increase in the expression of *PR10* and of the gene encoding phenylalanine lyase (*PAL)*, the key enzyme of salicylic acid biosynthesis in barley. The activation of defense gene expression in oeW1 plants coincided with an enhanced resistance towards powdery mildew, which in barley is independent of salicylic acid. Taken together, the results show that over-accumulation of WHIRLY1 in barley to levels of 10 or more, amplified the tradeoff between growth and stress resistance.

## INTRODUCTION

WHIRLY proteins are multifunctional DNA/RNA binding proteins localized to the DNA-containing organelles and the nucleus of higher plants (Krupinska *et al*., 2022). Investigations with mutants and knockdown plants have shown that WHIRLIES affect developmental processes and stress tolerance (Krupinska *et al*., 2022).

Initially, the WHIRLY1 protein has been identified as a transcriptional activator of the pathogen response gene *PR10a* in potato (Desveaux *et al*., 2000). Its binding to the promoters of PR genes that are enriched in elicitor response elements (ERE) was shown to depend on salicylic acid (SA) (Desveaux *et al*., 2004). In recent years it has been shown that the role of SA is not limited to pathogen defense but that SA has an essential role in the regulation of redox homeostasis and thereby affects plants’ responses towards abiotic and biotic stress (Mateo *et al*., 2006, Karpinski *et al*., 2013). Accordingly, the abundance of WHIRLY1 as a critical protein in SA signaling was shown to impact the plants’ tolerance to diverse abiotic stress situations as well as pathogen defense. In *whirly1 (why1)* Tilling mutants of Arabidopsis, in which the binding of WHIRLY1 to the promoter of *PR1* was reduced, resistance to *Peronospora parasitica* was relieved, too (Desveaux *et al*., 2004). Very recently, it has been reported that overexpression of *WHIRLY1* from *Vitis vinifera* under the control of a strong pathogen response promoter enhances resistance towards *Phytophthora capsica* (Lai *et al*., 2022).

Besides its positive effect on defense, WHIRLY proteins were also found to promote tolerance towards abiotic stress. In tomato plants, overexpression of *WHIRLY1* was shown to enhance thermotolerance by upregulating the expression of the *HSP21*.*5A* gene which has an ERE in its promoter and encodes an endoplasmic reticulum-localized heat shock protein (Zhuang *et al*., 2020a). Another study by the same research group showed that the plants overexpressing *SlWHIRLY1* had an enhanced chilling tolerance (Zhuang *et al*., 2019). Vice versa, tomato plants with an RNAi-mediated knockdown of *SlWHIRLY1* showed reduced resistance to chilling (Zhuang *et al*., 2019) and heat (Zhuang *et al*., 2020a). In maize and barley, it has been demonstrated that a reduction of WHIRLY1 negatively affects chloroplast development (Prikryl *et al*., 2008, Krupinska *et al*., 2019). Furthermore, barley plants deficient in WHIRLY1 were shown to be compromised in light acclimation (Saeid Nia *et al*., 2022).

Intriguingly, WHIRLY1 in barley was shown to locate in both, chloroplasts and nucleus of the same cell (Grabowski *et al*., 2008). In transplastomic tobacco plants synthesizing a tagged WHIRLY1 protein, this tagged protein was found in the nucleus indicating a translocation of WHIRLY1 from chloroplasts to the nucleus. In these plants, the expression of PR genes was enhanced (Isemer *et al*., 2012b). It has been hypothesized that storage of a transferable resistance protein such as WHIRLY1 in plastids might allow the plants to react immediately to pathogen attack avoiding the time and costs of gene expression (Krause and Krupinska, 2009). The translocation was suggested to occur in response to stress-associated redox changes in the photosynthetic apparatus (Foyer *et al*., 2014). WHIRLY1 is a positive regulator of both plant development and stress tolerance. Hence, WHIRLY1 promotes two traits that usually are inversely correlated. Indeed, enhanced stress tolerance coincides with lower growth and productivity (Herms and Mattson, 1992). This tradeoff is thought to be caused by resource restrictions demanding a prioritization of either growth or defense in response to environmental factors (Huot *et al*., 2014). The tradeoff is seemingly inevitable because the energy required for resistance is no longer available for biomass accumulation and production of seeds (Karasov *et al*., 2017). Thousands of genes are typically activated to fight a pathogen or cope with another stressful situation. Among others, the tradeoff between growth and defense is regulated by crosstalk between defense and growth hormones (Huot *et al*., 2014). Regulation of the level of free auxin is a significant determinant of adaptive growth in response to biotic and abiotic stress (Park *et al*., 2007). Recently, it has been demonstrated that MAP kinases activated during the immune response are involved in the downregulation of the expression of photosynthesis-associated genes, thereby exerting a negative impact on growth (Su *et al*., 2018).

Several studies with different dicot species have clearly shown a positive impact of over-accumulating WHIRLY1 on stress tolerance, however, without reporting effects on development and growth in these plants. Regarding the multifunctionality of WHIRLIES (Krupinska *et al*., 2022a) it is expected that other physiological parameters are altered besides stress tolerance. This study aimed to investigate the impact of a much higher abundance of multifunctional WHIRLY1 on plant growth and stress tolerance.

## RESULTS

### Overexpression of *HvWHIRLY1* altered the abundance of HvWHIRLY1 in chloroplasts and the nucleus

By transforming barley with *HvWHIRLY1* under the control of the constitutive *UBIQUITIN 1* promoter of maize (Figure 1a), three homozygous lines were selected and used for characterization. Immunoblot analysis with the HvWHIRLY1 specific antibody (Grabowski *et al*., 2008) revealed that in primary foliage leaves of line oeW1-14, the level of HvWHIRLY1 was enhanced by a factor of about 50 (Figure 1b) as it was also in line oeW1-2 (Figure S1a). For comparison, in line oeW1-15 the level of WHIRLY1 was enhanced by a factor of 10 (Figure 1b).

**Figure 1.**
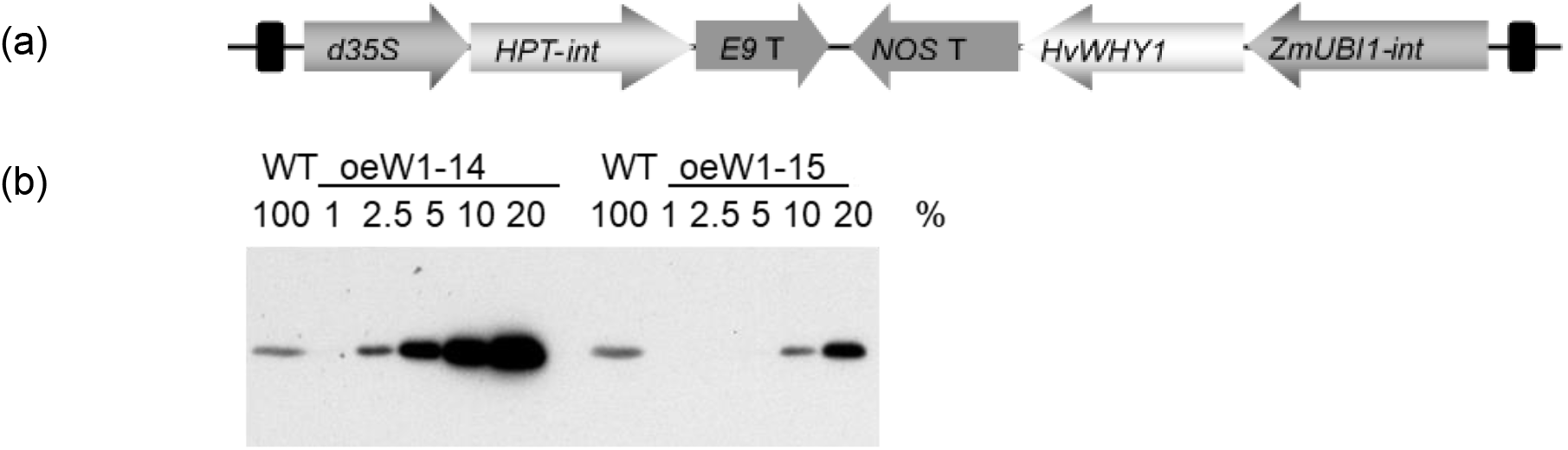
Overexpression of *HvWHIRLY1* in barley. (a) Schematic drawing of the T-DNA used for overexpression of *HvWHIRLY1* under control of the *UBIQUITIN 1* promoter of maize. (b) Accumulation of the WHIRLY1 protein in total protein extracts derived from primary foliage leaves of the two transgenic lines, oeWHIRLY1-14 and oeWHIRLY1-15. For comparison, different amounts of leaf protein were used and indicated as a percentage of protein from wild-type plants (WT). d35S – doubled enhanced *CaMV 35S* promoter, HPT-int – *Hygromycin phosphotransferase* gene with *StLS1* intron, *E9* T - Terminator of *Rbcs-E9* gene, ZmUBI1-int – maize *UBIQUITIN 1* promoter with the first intron, *HvWHY1* – barley *WHIRLY1, NOS* T *– Agrobacterium tumefaciens Nopaline synthase* gene termination signal.

WHIRLY1 is dually located in chloroplasts and nucleus. To investigate whether the over-accumulation of the protein occurred in both compartments and whether the relative distribution between chloroplasts and nucleus is altered by the overexpression of *WHIRLY1* leaves (Fig. 2a) WHIRLY1 abundance was immunologically investigated in chloroplast and nuclei fractions prepared from primary foliage leaves of the oeW1-15 and the oeW1-14 lines. Theoretically, excess WHIRLY1 could accumulate only in chloroplasts or could also accumulate in nuclei, whereby the ratio between chloroplast and nuclear WHIRLY1 could be similar to in WT plants or could be shifted towards the nuclear form (Figure 2a). Immunoblots with the specific antibody for HvWHIRLY1 showed that the abundance of HvWHIRLY1 was enhanced in both chloroplasts and nuclei (Figure 2b, Figure S1b). Thereby the relative increase in quantity in both lines was higher in nuclei than in chloroplasts (Fig. 2c). Considering that the proteins in chloroplasts and nucleus have the same molecular weight, the higher abundance in the nucleus likely results from an enhanced flux of protein from chloroplasts to the nucleus. This result suggests that the capacity to sequester HvWHIRLY1 in chloroplasts is saturated, and relatively more WHIRLY1 is transferred to the nucleus (Figure 2).

**Figure 2.**
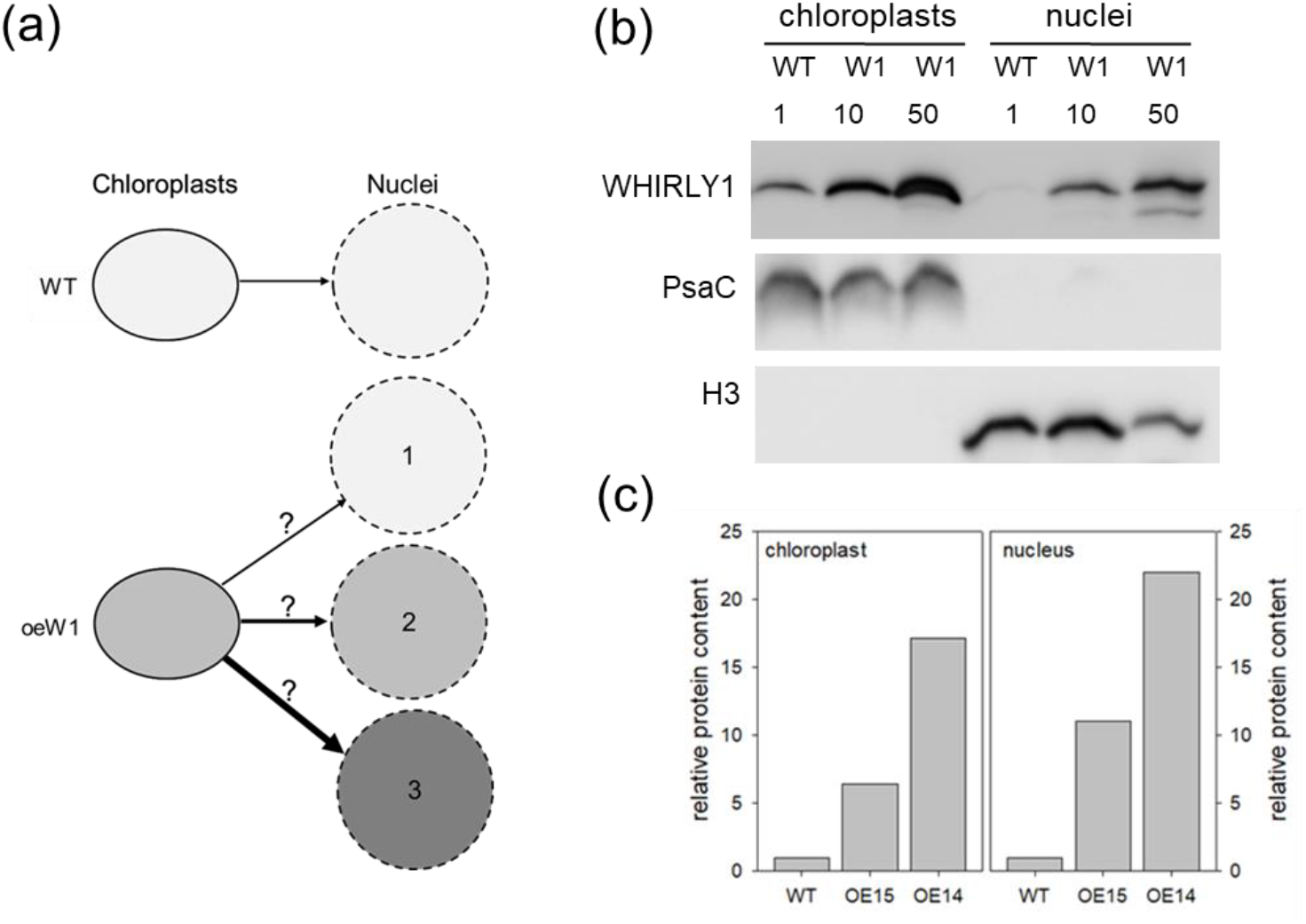
Relative WHIRLY1 levels in extracts from chloroplasts and nuclei, respectively. (a) Putative consequences of *WHIRLY1* overexpression (oeW1) for the distribution of the WHIRLY1 protein between chloroplasts and the nucleus. Different levels of WHIRLY1 are illustrated by light, medium, or dark grey symbols that represent chloroplasts or nuclei. The distribution in chloroplasts and nuclei of the wild type (WT) is normalized (light grey area). Either the transfer of WHIRLY1 to the nucleus will be not altered in comparison to the wild type (1), or the transfer will be enhanced resulting in a similar relative over-accumulation of WHIRLY1 in chloroplasts and the nucleus (2) or the transfer will be relatively higher as expected resulting in a higher relative abundance of WHIRLY1 in the nucleus compared to chloroplasts (3). (b) Subcellular fractions were prepared from primary foliage leaves of WT, oeW1-15 (10-fold accumulation of WHIRLY1), and oeW1-14 (50-fold accumulation of WHIRLY1) plants. Immunoblot analysis was performed with extracts from chloroplasts (CP) and nuclei (N). Each lane was loaded with 6 μg of protein. To show the purity of fractions, antibodies directed towards PsaC and histone 3 (H3) have been used. (c) Relative abundances of the WHIRLY1 protein were calculated from the signal intensities measured by the ChemiDoc MP Imaging Systems after different times of exposure using the Image Lab 6.1 software. The relative intensities of the WHIRLY1 signals detected in chloroplast and nuclei samples, respectively, are based on the signal intensities of WT samples.

### Growth of barley oeW1 plants

To investigate whether WHIRLY1 overaccumulation has consequences for growth, the lengths of primary foliage leaves were measured every day until they were fully expanded (Figure 3). An apparent growth reduction correlated with an abundance of WHIRLY1. Growth curves showed that the seedlings of oeW1-10 were longer than those of the oeW1-50 line (Figure 3). The growth kinetics did not differ between WT and oeW1 seedlings. The maximal expansion of the primary foliage leaves was terminated at the same time after sowing, i.e. at 9 days (Fig. 2a).

**Figure 3.**
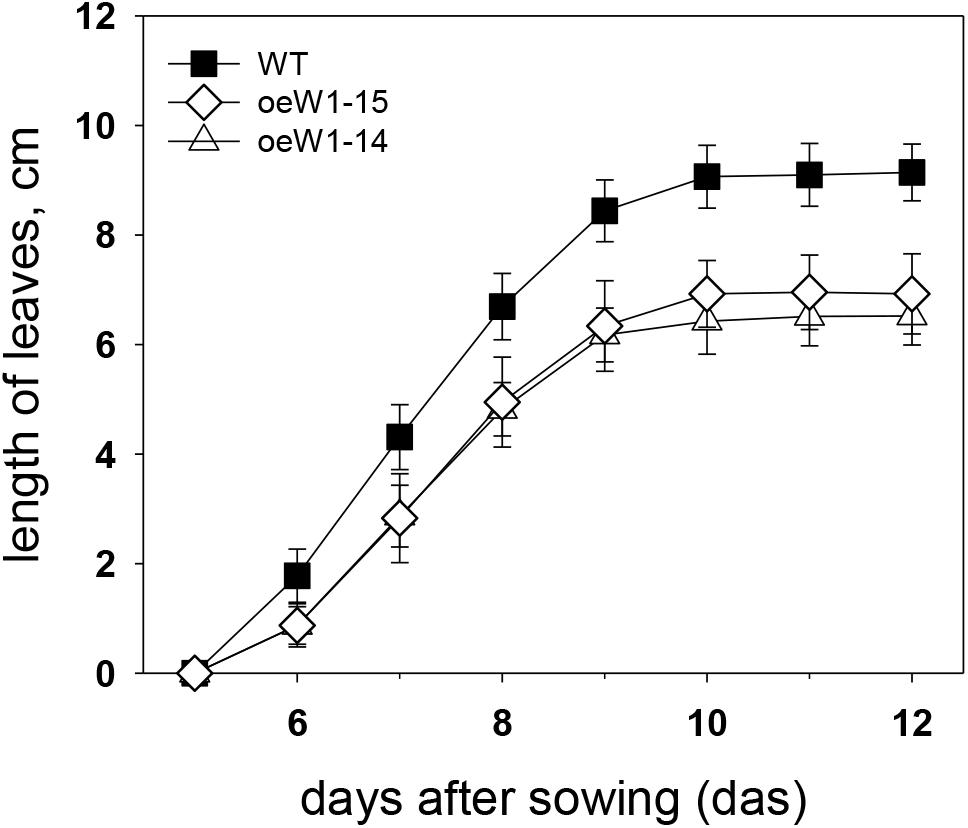
The lengths of entire primary foliage leaves of WT, oeW15, and oeW14 were determined by measuring the lengths from the kernel, where the leaf sheath starts, to the leaf tip at 5-12 days after sowing. Symbols represent means ± SD of n=13-20 leaves.

### Characterization of the photosynthetic apparatus

To investigate whether changes in the photosynthetic apparatus are responsible for the reduced growth of the oeW1 seedlings, the functionalities of the two photosystems were examined during the development of barley seedlings in a daily light/dark regime. Chlorophyll fluorescence measurements revealed that the maximal quantum yield of photosystem II, F_V_/F_M,_ which is a measure of photosystem II efficiency, in WT seedlings was already relatively high at 7 das (0.7) and increased to nearly 0.8 at 10 das. In comparison, F_V_/F_M_ of the oeW1-50 leaves stayed rather low, reaching a maximal value of about 0.5 at 10 das (Figure 4a).

**Figure 4.**
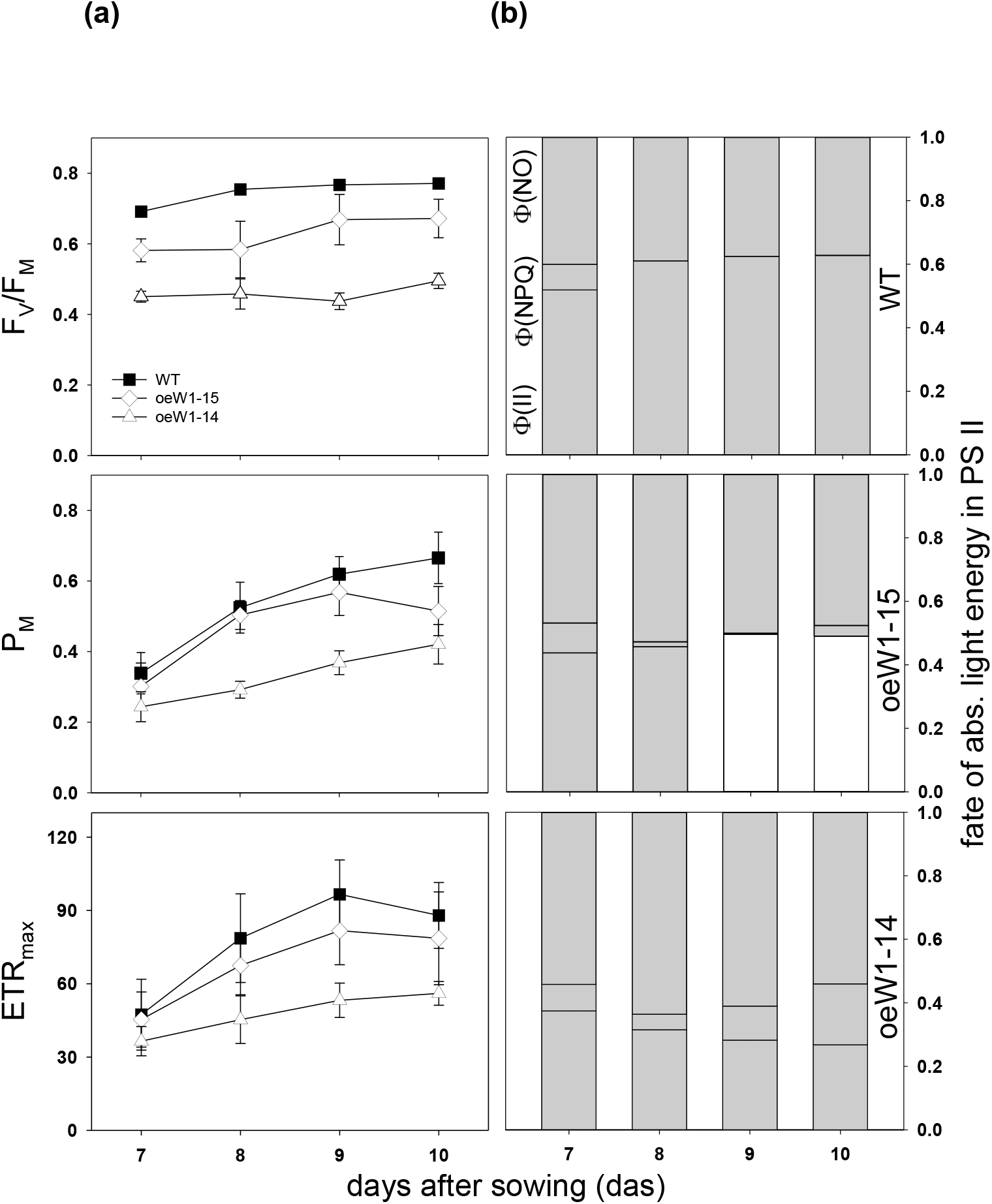
Characterizing the photosynthetic apparatus in primary foliage leaves of WT, oeW1-14, and oeW1-15 seedlings grown under a daily light-dark cycle (L/D). (a) The optimal quantum yield of photosystem II, F_V_/F_M_, the maximum P700 signal (P_M_), and the maximal electron transport rate (ETR_max_) were measured at room temperature at different days after sowing (7-10 das). Depicted values are mean ± SD of n=6 leaves. (b) Fate of the light energy absorbed at PS II was determined at an irradiance of 60 μmol m^-2^ s^-1^. The quantum yield of photochemistry (Ф(II)), of regulated non-photochemical quenching (Ф(NPQ)) and non-regulated non-photochemical quenching (Ф(NO)) were calculated according to the formulas given in Materials and Methods. Columns are means ± SD of n=6 leaves.

For comparison, in oeW1-10 seedlings, F_V_/F_M_ had a value of 0.6 at 7 das and a value of 0.7 at 9 and 10 das (Fig. 4a). In contrast to F_V_/F_M,_ which barely changed in wild-type leaves during the growth period investigated, the capacity of photosystem I measured by the maximal absorbance change of P_700_ (P_M_) increased from 0.3 at 7 das until 0.7 at 10 das (Figure 4a). The values stayed significantly lower in both oeW1 lines, whereby oeW1-14 seedlings had lower values than oeW1-15 seedlings (Fig. 4a).

In addition to the efficiency of photosystem II, the maximum electron transport rate of photosystem II (ETR_max_) was reduced in the primary foliage leaves of oeW1 seedlings measured on different days after sowing (7-10 das). The results showed that oeW1-14 leaves had only about 50% of the electron transport capacity of WT leaves, while oeW1-15 leaves had about 80% of the WT level (Figure 4a). Whereas the transport rate in WT and oeW1-15 leaves was maximal already at 9 das, it still increased in oeW1-14 leaves from 9 das until 10 das. An analysis of the partitioning of absorbed energy in photosystem II revealed that the quantum yield of photosystem II (Ф(II)) was lower in oeW1 plants at all stages of development compared to WT plants. The remaining fraction was dissipated mainly as heat or fluorescence (Ф(NO)). Only a small fraction was used for non radiative dissipation in the oeW1 leaves (Figure 4b).

To investigate putative differences in the composition of the photosynthetic apparatus, the concentrations of pigments and the relative abundances of photosynthesis-associated proteins were analyzed. Pigment analyses by HPLC showed that the chlorophyll content per leaf area of primary foliage leaves from seedlings of the oeW1-50 leaves was lower than the chlorophyll contents of line oeW1-15 and the wild-type, which were similar (Figure 5). The ratio of chlorophyll a/b was reduced in both oeW1 lines (Figure 5) and was independent of the leaves developmental stage.

**Figure 5.**
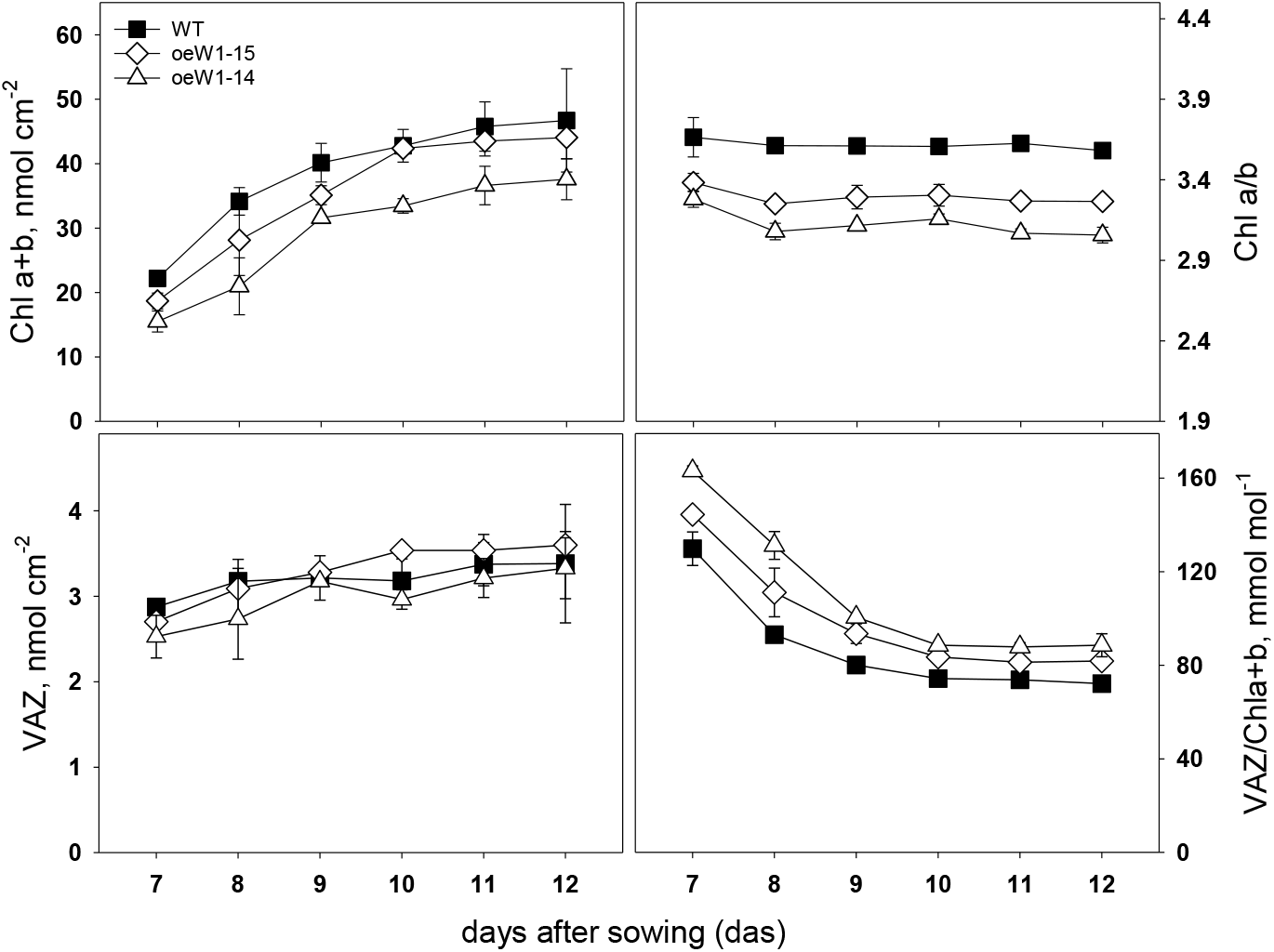
Pigment content of primary foliage leaves of barley wild type, oeW1-14 and oeW1-15 lines. Leaves were collected in the growth period from day 7 until day 12 after sowing. Depicted values are means±SD of n=3 leaves.

The content of xanthophyll cycle pigments was similar in all genotypes. However, the xanthophyll pool/chlorophyll ratio was higher in the leaves of the oeW1 lines, in particular in the oeW1-50 leaves.

Protein extracts from primary foliage leaves of WT and oeW1 seedlings were immunologically analyzed for the levels of central photosystem I (PsaA), photosystem II (D1/PsbA) proteins, and two light-harvesting proteins, i.e. LHCA1 and LHCB1, respectively. As already reported, the abundance of WHIRLY1 in the WT declined with the increasing age of the leaves (Kucharewicz *et al*., 2017, Krupinska *et al*., 2019). The levels of all tested proteins were similar in the WT and the oeW1 lines. (Figure 6). The staining also indicates that the two subunits of RubisCO had the same abundance as in the WT and that the levels are stable during development from 7 das until 12 das.

**Figure 6.**
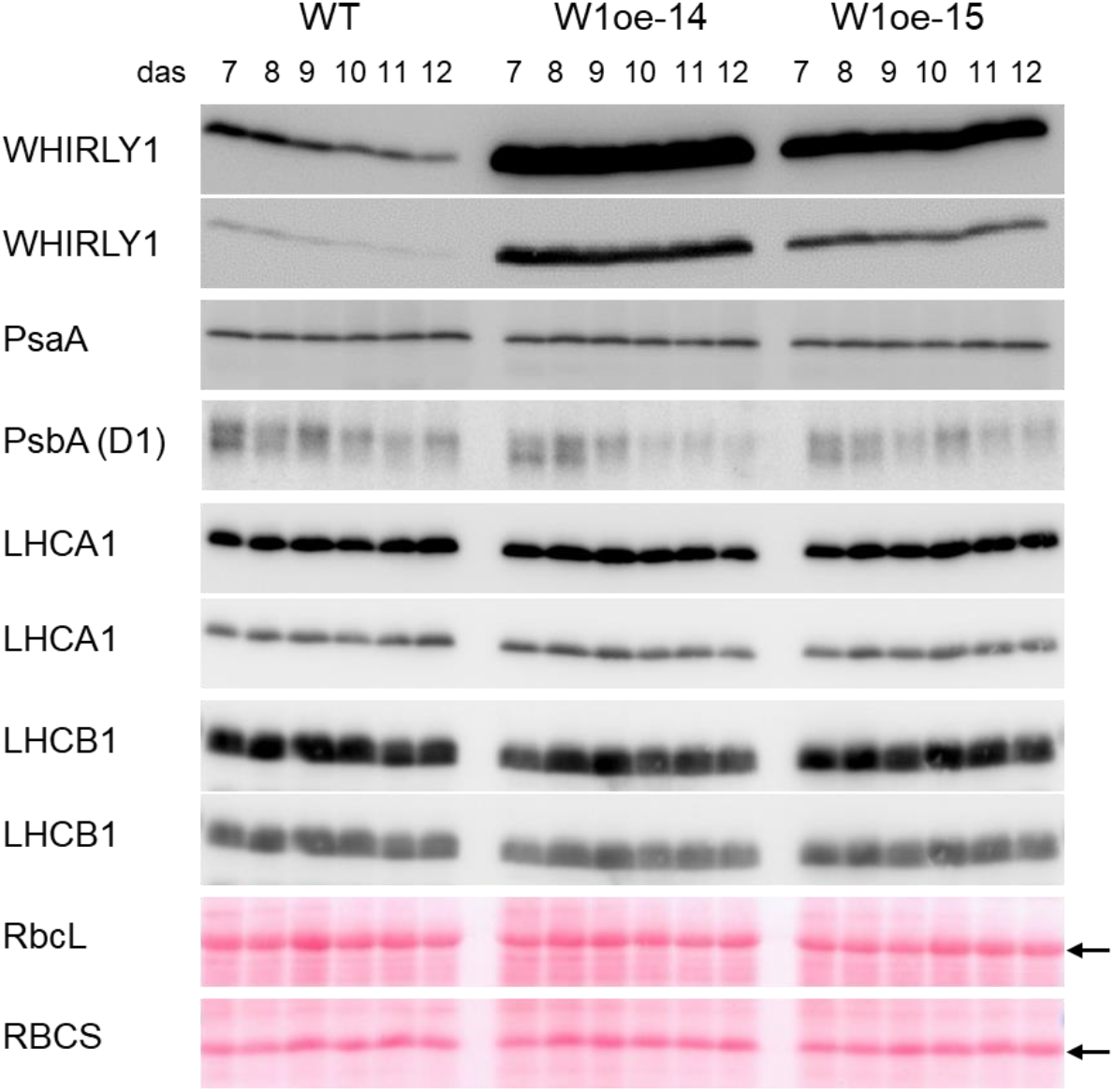
Relative amounts of proteins of the photosynthetic apparatus in primary foliage leaves of the WT,the oeW1-14 line, and the oeW1-15 line. Protein extracts were prepared from the leaves at different times after sowing (7-12 das). Immunological analyses were performed with specific antibodies directed towards WHIRLY1 and selected proteins of the photosynthetic apparatus: PsaA, PsbA (D1), LHCA1, and LHCB1. In the case of WHIRLY1, LHCA1 and LHCB1, two different exposures are shown, respectively. At the bottom, Ponceau stained gel parts showing the large and the small subunits of RubisCO (indicated by arrows) are presented.

In order to investigate gene expression in chloroplasts from oeW1 leaves in comparison to WT leaves, mRNA levels of genes encoding central components of the photosynthetic apparatus were analyzed by RT-PCR (Figure S2). While mRNA levels of all genes declined during the development of WT leaves, the mRNAs stayed at relatively high levels during the development of the oeW1 leaves. While in RNAi-W1 plants, plastid gene expression was mainly due to the activity of the nuclear-encoded RNA polymerase (NEP) (Krupinska *et al*., 2019), in the oeW1 lines transcripts of both NEP (*rpoB, clpP*) and PEP (*psbE*) were present at higher levels than in WT plants. This result indicates that overexpression of *WHIRLY1* did not hamper transcription in chloroplasts.

### Chloroplast ultrastructure and nucleoid morphology

When primary foliage leaves of barley were fully expanded (10 das), ultrathin sections from WT and oeW1-14 seedlings grown in a daily light/dark cycle were fixed for ultrastructural analyses by transmission electron microscopy. While mitochondria and peroxisomes looked rather similar in WT and oeW1 samples (Figure 7a), chloroplasts showed noticeable morphological differences (Figure 7b). Chloroplasts of oeW1 plants apparently contained more plastoglobules (Figure 7). Plastoglobules are lipoprotein particles surrounded by a lipid monolayer, which is contiguous with the outer leaflet of thylakoid membranes. They contain mainly isoprenoid-derived lipophilic compounds and function in remodeling the lipid phase of thylakoids (van Wijk and Kessler, 2017). An increase in the number and/or size of plastoglobules was reported to indicate excess light in the photosynthetic apparatus (Brehelin *et al*., 2007, Rottet *et al*., 2016) and was observed during various stressful situations (Lichtenthaler, 2013, van Wijk and Kessler, 2017). Thylakoids in the chloroplasts of oeW1 leaves showed a tendency to swell. Following the lower photosynthetic activity of oeW1 leaves (Figure 3), chloroplasts of the oeW1 plants did not contain starch grains which were frequently observed in the wild-type chloroplasts.

**Figure 7.**
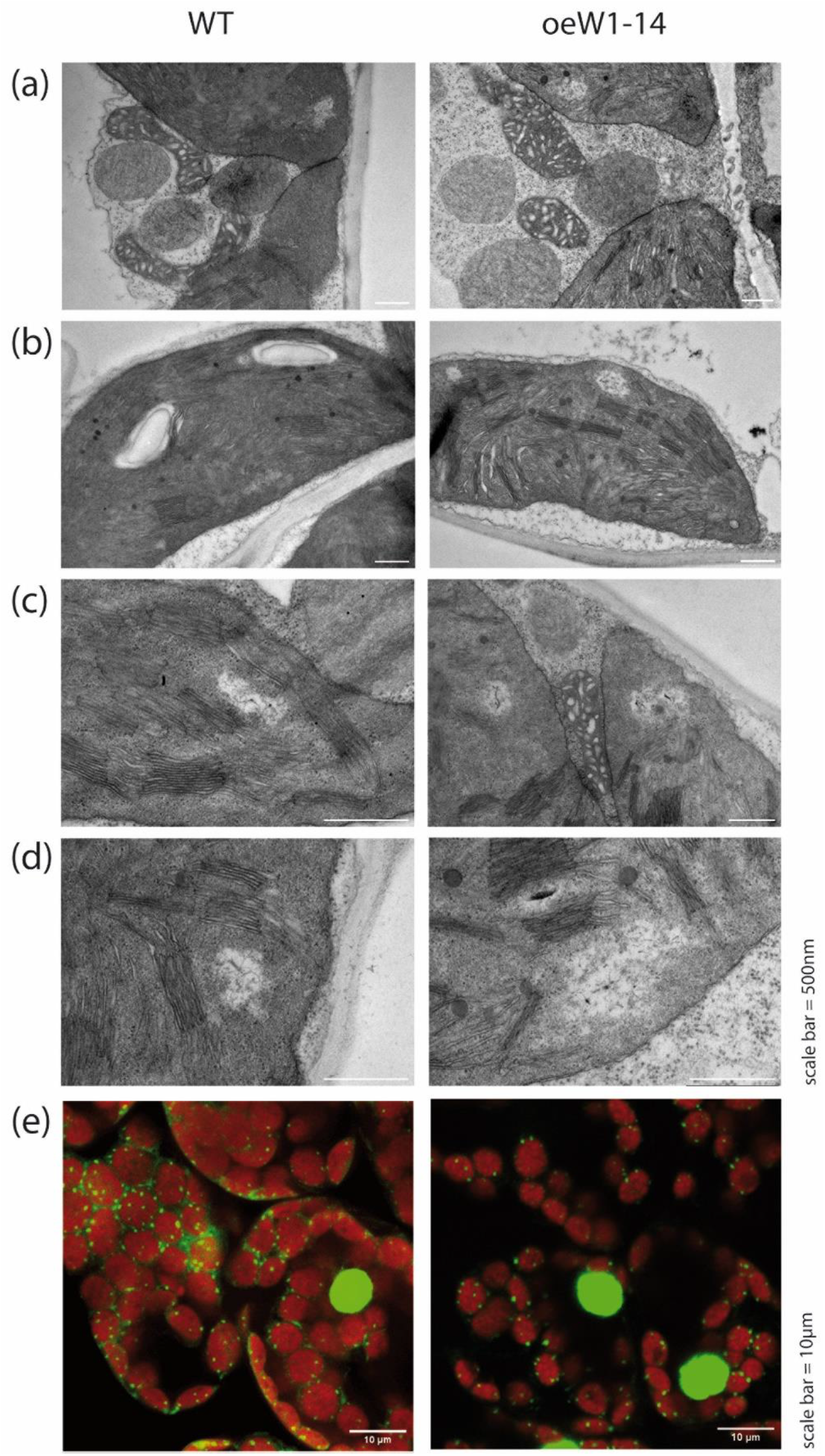
Microscopic analyses of chloroplasts and nucleoids from WT and oeW1-14 seedlings by transmission electron microscopy (a-d) and confocal fluorescence microscopy (e) where the green fluorescence was emitted by DNA stained with SYBR Green. Samples were excited by an argon laser at 488 nm (5% power). Emission was detected between 510-570 nm (HV750) and 690-760 nm (HV480).

Considering that WHIRLY1 is a major nucleoid-associated protein (Pfalz *et al*., 2006, Krupinska *et al*., 2022b), special attention was committed to the structure of nucleoids in mature chloroplasts of the primary foliage leaves of WT and oeW1 seedlings. Ultrastructural analyses did, however, not reveal apparent differences between nucleoids in WT and oeW1 sections (Fig. 6c-d). In addition, nucleoids of mature chloroplasts were also visualized by light microscopy after staining sections with SYBR Green (Figure 7e). By this procedure, neither differences in size nor the distribution of nucleoids were observed between the two genotypes.

Previous studies revealed a profound impact of WHIRLY1 on the packaging of plastid DNA (Krupinska *et al*., 2014) and bacterial nucleoids (Oetke *et al*., 2022). Considering that WHIRLY1 also plays a significant role in chloroplast development (Prikryl *et al*., 2008, Krupinska *et al*., 2019, Krupinska *et al*., 2022), it might be possible that putative differences in nucleoid morphology depend on the developmental stage of plastids. Using the developmental gradient of the leaves of small-grained cereals (Boffey *et al*., 1980), putative development-related changes in nucleoid morphology were investigated by staining sections prepared from the base and the middle part of the leaves of WT, oeW1-2 having a 50-fold over-accumulation as in oeW1-14 plants, and oeW1-15 seedlings with a 10-fold over-accumulation of WHIRLY1, respectively. Although WHIRLY1 abundances in chloroplasts are dramatically different between WT and an oeW1-50 line, no apparent differences in nucleoid morphology were observed. In all genotypes, nucleoids in undifferentiated plastids at the base are arranged like pearls on a string. In contrast, nucleoids in mature chloroplasts are dispersed inside the chloroplasts (Figure S3) due to their attachment to thylakoid membranes (Powikrowska *et al*., 2014).

### Hormone levels and defense-related gene expression

To elucidate whether the reduced growth of the oeW1 plants was related to changes in the levels of growth hormones, cytokinins and auxins were determined at 7 and 10 das in primary foliage leaves of plants grown at a light intensity of 100 μmol m^-2^ s^-1^. These measurements revealed that independent of the developmental stage, the levels of the cytokinin N^6-^ isopentenyl adenosine (iPR) were reduced by about 30% or 60% in the primary foliage leaves of oeW1-15 or oeW1-14 plants, respectively (Figure 8). For comparison, indole-3-acetic acid (IAA) levels were similar among the lines. At 10 das a reduction in the level of IAA by about 20% was measured in the leaves of the oeW1-14 line (Figure 8).

**Figure 8.**
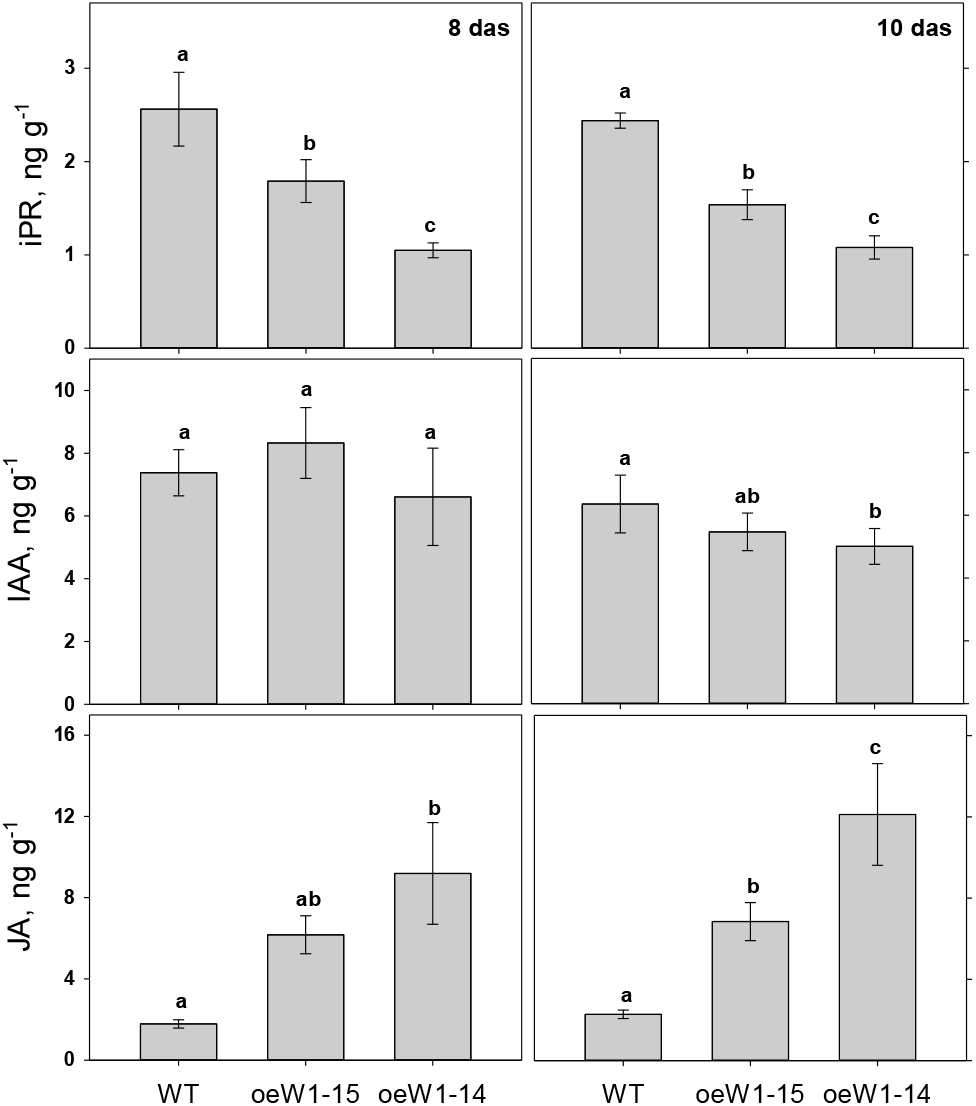
Hormone levels in primary foliage leaves of the WT in comparison to the oeW1-15 and oeW1-14 lines: N^6^-isophentenyladenosine (iPR), indole acetic acid (IAA) and jasmonic acid (JA). Leaves were collected at 8 das and at 10 das, respectively. Columns are means ± SD of n=5 leaves.

For comparison, the levels of hormones involved in defense were determined. While free salicylic acid was too low to be determined, its catabolite and storage compound were detectable. It has been shown that in barley during pathogen defense, SA is not produced via the isochorismate (ICS) pathway (Vlot *et al*., 2009, Rekhter *et al*., 2019), but rather by the phenylpropanoid pathway controlled by phenylalanine lyase (PAL) (Qin *et al*., 2019). The levels of 3,4-dihydroxy benzoic acid were about twofold higher in young leaves of the oeW1-14 leaves compared to the other lines, but this difference disappeared when leaves were collected at 10 das (Figure S6). Levels of SA glucosides which is a storage form of SA (Vlot *et al*., 2009), showed a tendency to be higher in the leaves from oeW1-14 plants, in particular in young leaves (7 das) (Figure S6). The determination of jasmonic acid (JA) revealed that at 8 das overexpression of *HvWHIRLY1* significantly increased its level by a factor of 3 or 5 in oeW1-15 or oeW1-14 lines, respectively. Thereby the basis level measured in the WT leaves was slightly enhanced at 10 das, compared to 8 das (Figure 8). Taken together, the results revealed that over-accumulation of WHIRLY1 induced reciprocal changes in the levels of iPR and JA which might cause a shift from growth to defense.

To investigate whether according changes in gene expression accompanied the transition from development to defense, mRNA levels of key enzymes in the biosynthesis of defense hormones were determined besides the levels of *WHIRLY1* mRNA and *PR1* mRNA by quantitative real-time PCR. The result showed that *HvWHIRLY1* had an up to 20-fold higher mRNA level in primary foliage leaves of oeW1-14 seedlings compared to the WT. *PR1* is known as a marker of SAR (Linthorst, 1991). While in Arabidopsis and other dicots, it was reported to be a target gene of salicylic acid (Van Loon and Van Strien, 1999, Golshani *et al*., 2015), *PR1* in rice was shown to accumulate in response to JA (Rakwal and Komatsu, 2000, Jwa *et al*., 2006). In this study, barley *PR1* expression was upregulated in WT and oeW1 seedlings when the leaves became fully expanded. While in the WT, *PR1* expression was upregulated by a factor of 6 at 10 das and by a factor of 11 at 12 das, in oeW1 seedlings, expression of the gene was highly upregulated by a factor of 120 at 10 das (Figure 9). Upregulation of *PR1* in the oeW1 seedlings was neither accompanied by upregulation of the gene encoding PAL nor ICS, which are the key enzymes of the two pathways of salicylic acid biosynthesis (Vlot *et al*., 2009) (Figure 9).

**Figure 9.**
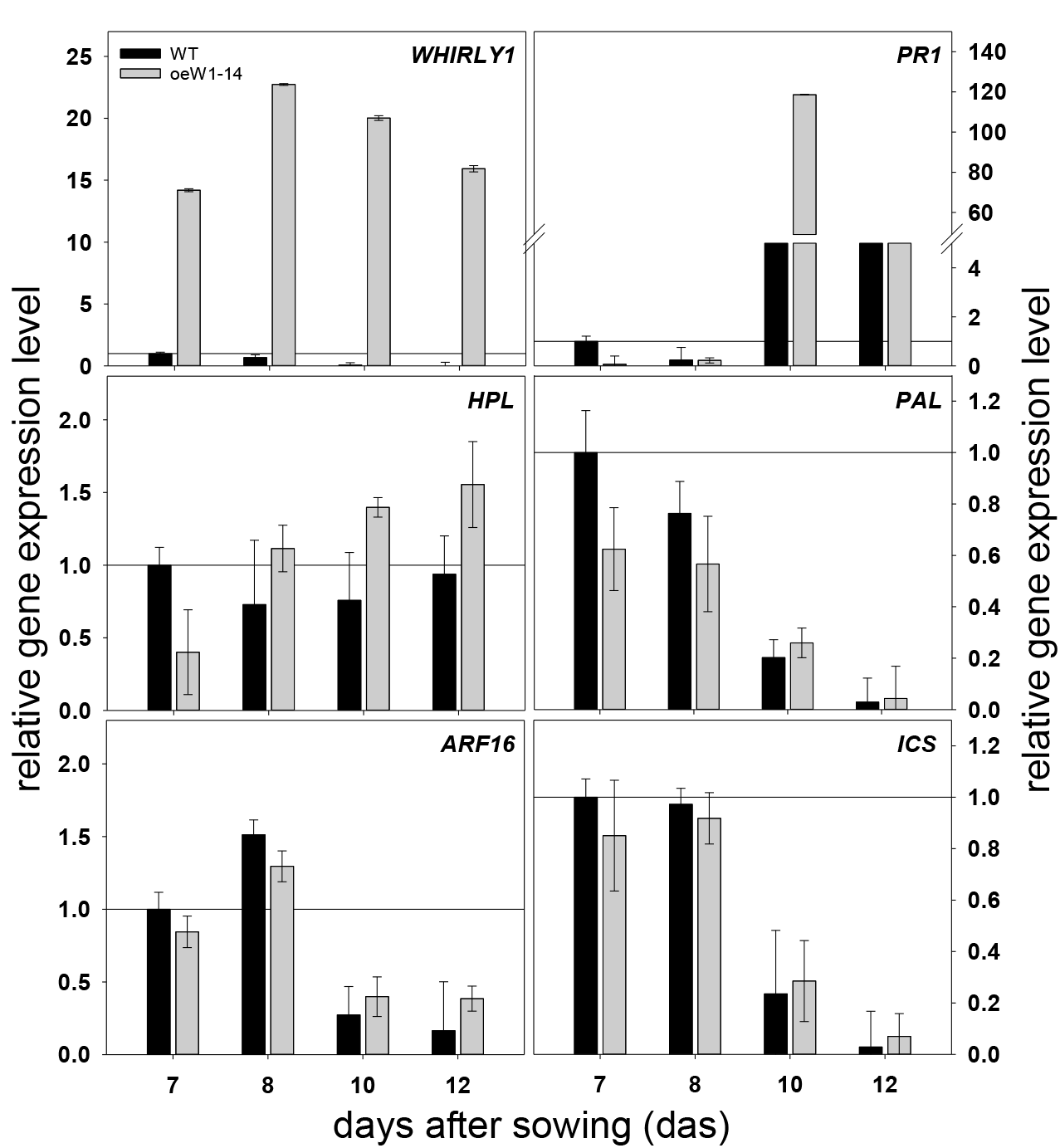
Expression of *HvWHIRLY1* and selected stress-associated genes measured by qRT-PCR using *GAPDH* (see Material and Methods) as standard. RNA was extracted from primary foliage leaves of WT and oeW1-14 seedlings grown for different times (7, 8, 10 and 12 das) in a daily light/dark cycle. RNA levels were compared to those of the wild type at 7 das that were set to 1 and represented by horizontal lines. Columns are means ± SD of n=3 samples (each sample comprised 10 pooled leaves).

Expression of the general stress associated *HPL* gene encoding hydroperoxide lyase, a chloroplast protein of the oxylipin pathway shown to protect against photoinhibition (Savchenko *et al*., 2017), was only upregulated by about 50% in fully expanded primary foliage leaves of the oeW1 seedlings. This gene was chosen because it is known to be a stress indicator gene regulated by retrograde signaling during stress in Arabidopsis (Xiao *et al*., 2012, Xiao *et al*., 2013). Its expression barely changed in WT seedlings during normal growth (Figure 9). In contrast to *PR1*, the expression of *THIO1*, another barley defense gene (Leybourne *et al*., 2022), was downregulated during growth in both WT and oeW1 seedlings (Figure S4).

### Response of oeW1 plants to high light

In Arabidopsis, defense signaling is also activated in response to high light (Mateo *et al*., 2006, Karpinski *et al*., 2013). To induce high light stress, oeW1-14 and WT seedlings were grown in continuous light of 350 μmol m^-2^ s^-1^ (HL) and were compared to seedlings grown at only 100 μmol m^-2^ s^-1^ (LL) as described previously (Swida-Barteczka *et al*., 2018). Growth at high light leads to a decrease in the chlorophyll content of both WT and oeW1 plants. The reduction in chlorophyll content of oeW1 plants was significantly more pronounced than the WT (Figure 10a) but was not as prominent as in the case of the *WHIRLY1* knockdown plants prepared by RNAi (Swida-Barteczka *et al*., 2018). In WT seedlings, F_V_/F_M_ was not affected by higher irradiance during growth, while it even slightly increased in the case of oeW1-14 seedlings (Figure 10a).

**Figure 10.**
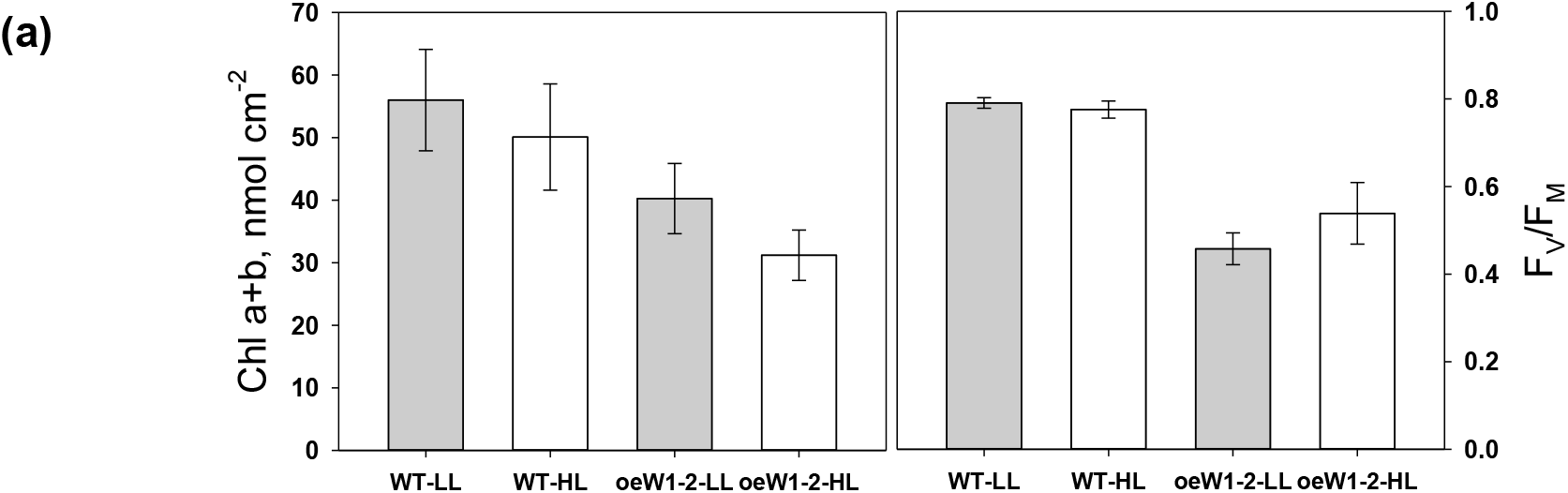
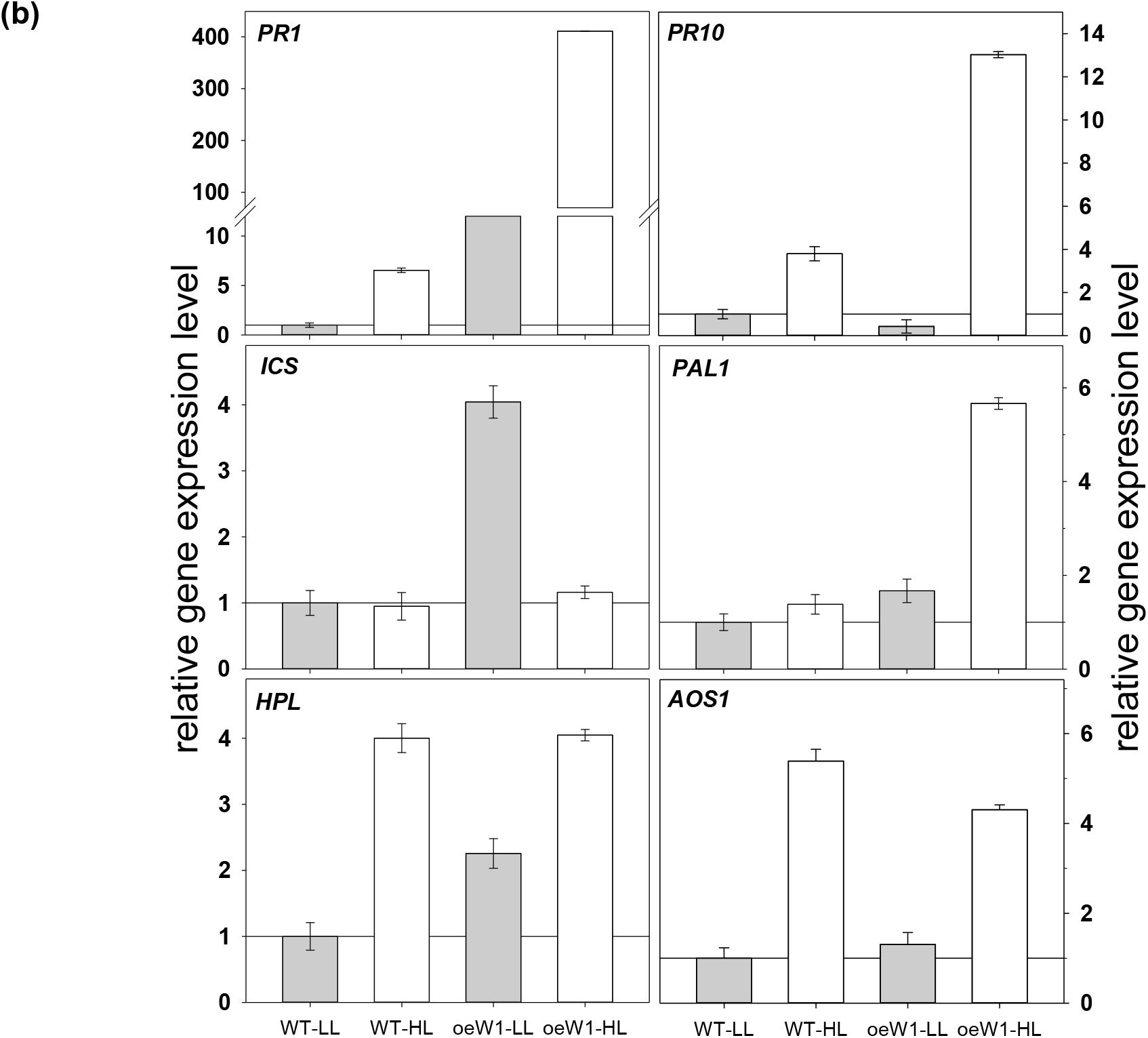
Characterization of primary foliage leaves of WT and oeW1-2 seedlings during growth in continuous light of different irradiance (low ligh=LL in grey and high light =HL in open columns). (a) Chlorophyll content and F_V_/F_M,_ (b) expression of defense-related genes putatively associated with HL stress and SA: *PR1, PR10, ICS, PAL, HPL, PAL1, AOS*. mRNA levels were measured by qRT-PCR using *GAPDH* as standard gene. The levels were compared to those of the wild type grown at LL that were set to 1 and represented by horizontal lines.

When WT seedlings were grown at HL, *PR1*, and *PR10* expression levels were elevated compared to the levels determined in LL-grown plants. This result follows the idea that SA is involved in response to HL. Overexpression of *WHIRLY1* led to a dramatic increase in the expression of both PR genes (Figure 10b). Moreover, over-accumulation of HvWHIRLY1 led to enhanced expression of *PAL*, which was more pronounced at HL than at LL (4-fold in comparison to LL) (Figure 10b). Expression of *PAL* but not of *ICS* was also slightly enhanced in the WT at HL. In comparison, *ICS* expression was enhanced in oeW1 plants only at LL, but not at HL. The high expression of *ICS* at LL could be related to an increased demand for phylloquinone (Qin *et al*., 2019).

In addition, the expression of genes encoding two key enzymes of the two branches of the oxylipin biosynthesis in chloroplasts (Savchenko *et al*., 2017) was determined, i.e. HPL leading to the biosynthesis of aldehydes and allene oxide synthase (AOS), a key enzyme of JA biosynthesis (Delker *et al*., 2006). In the WT, *HPL* was upregulated by 4-fold, while AOS was upregulated by a factor of 5.5 (Figure 10b). In oeW1 seedlings, expression of *HPL* was already enhanced at LL and was only upregulated by 40% in HL compared to LL. Indeed, the expression levels at HL were identical between WT and oeW1 plants. The expression of *AOS* is upregulated likewise in HL in both the WT and the oeW1 plants.

### Response of oeW1 plants to powdery mildew

To investigate the impact of WHIRLY1 accumulation on pathogen resistance, leaves were inoculated with spores of the powdery mildew fungus *Blumeria graminis*, an important barley pathogen. The susceptibility to powdery mildew was compared among WT, oeW1-14, oeW1-2 (two lines over-accumulating WHIRLY1 by a factor of 50), and two barley plants with an RNAi-mediated knockdown of *HvWHIRLY1*, W1-1 and W1-7 (with 10% and 1% of the protein in WT, respectively), which had been used in several investigations before (Krupinska *et al*., 2014b, Krupinska *et al*., 2019). Both oeW1-14, oeW1-2 were less susceptible to powdery mildew than the WT, as determined by estimating the percentage of the leaf surface infected by the fungus (Figure 11). Inversely, the leaves of the *WHIRLY* knockdown plants (W1-1 and W1-7) were more susceptible to inoculation with powdery mildew spores. The results show that a high abundance of WHIRLY positively affects the resistance towards powdery mildew.

**Figure 11.**
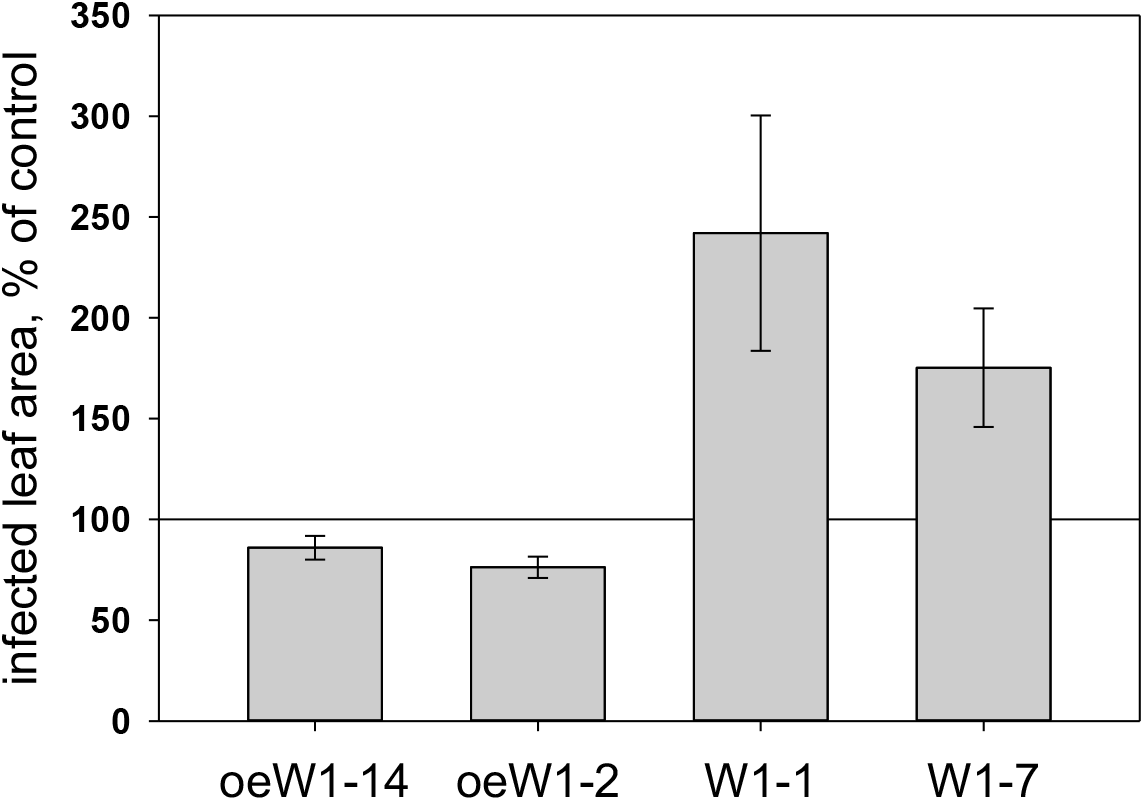
Infection of barley leaves by powdery mildew (*Blumeria graminis*). WT leaves were compared with leaves of two oeW1 lines over-accumulating WHIRLY1 by a factor of 50 (oeW1-14, oeW1-2), and with the *knockdown* plants having residual amounts of about 10% (W1-1) or 1% (W1-7) of WHIRLY1 protein (Krupinska *et al*., 2014b). Susceptibility was determined by the percentage of leaf area infected by the fungus. The susceptibility of the WT has been defined as 100% represented by horizontal lines.

## DISCUSSION

Overexpression of *WHIRLY1* in barley resulted in an up to 50-fold higher abundance of the WHIRLY1 protein, an improved tolerance towards powdery mildew, and diminished growth, indicating a typical tradeoff between growth and defense (Herms and Mattson, 1992, Huot *et al*., 2014). Although the tradeoff has often been explained by the competition of energy requirements of defense responses in relation to those for growth and reproduction, this apparently plausible explanation has also been questioned. Instead, the dilemma between development and defense was shown to stem from antagonistic crosstalks between growth and defense-related hormones (Karasov *et al*., 2017), which can be uncoupled in mutants (Campos *et al*., 2016).

### Impact of *WHIRLY1* overexpression on growth and photosynthesis

For oe*AtWHIRLY1* plants, no obvious phenotype has been reported (Isemer *et al*., 2012a). A more detailed characterization has been performed with tomato lines overexpressing *SlWHIRLY1* (Zhuang *et al*., 2019). In these plants, the mRNA level increased dramatically by factors of about a thousand. In contrast, the protein level was only enhanced by an estimated factor of approximately five (estimated from Figure 2 in Zhuang *et al*. 2019). No significant difference was observed in the phenotypes between tomato oe*SlWHIRLY1* lines and the wild type at standard growth conditions. However, under chilling conditions, the oe*SlWHIRLY1* lines grew better than the wild-type (WT) coinciding with a reduced level of ROS, as shown by fluorescence after staining with H2DCFDA (Zhuang *et al*., 2019). At the ultrastructural level, the oe*SlWHIRLY1* plants were shown to retain intact grana thylakoids and to accumulate less starch in chilling conditions. However, in contrast to the barley lines, overexpressing *HvWHIRLY1* (oeW1), under control conditons the abundance of starch grains did apparently not differ between WT and oe*SlWHIRLY1* plants (Zhuang *et al*., 2019). Also in contrast to the barley oeW1 lines, the oe*SlWHIRLY1* plants showed no difference in F_V_/F_M_ at 25°C and even higher F_V_/F_M_ values under chilling conditions (Zhuang *et al*., 2020b). Also, in contrast to the barley oeW1 plants, RubisCO content was higher in the oe*SlWHIRLY1* plants than in the WT, both at 25°C and 4°C (Zhuang *et al*., 2020b). Under heat stress, the oe*SlWHIRLY1* plants showed less wilting than WT tomato plants coinciding with increased sugar content and a reduced level of ROS (Zhuang *et al*., 2020a).

In contrast to the barley oeW1 plants, the two *WHIRLY1* overexpressing dicot species investigated didn’t show a pronounced decrease in growth under standard conditions. Compared to the barley lines used in this study, over-accumulation of the protein in tomato is relatively low and could be a reason for the discrepancies between barley and tomato. Alternatively, the growth-related difference between WHIRLY1 over-accumulation in barley on the one hand, and tomato or Arabidopsis, on the other hand, could be due to differences in the impact of WHIRLY1 proteins on chloroplast nucleoid architecture. Only in monocots WHIRLY1 proteins were shown to have a specific PRAPP motif required for the compaction of nucleoids (Oetke *et al*., 2022). However, despite the over-accumulation of WHIRLY1, nucleoids did not show differences in their compactness and organization between WT and oeW1 plants as investigated by DNA staining (Figure 7, S3). This result is in line with the almost normal levels of plastid-encoded mRNAs (Figure S2) and the unvaried protein composition of the photosynthetic apparatus (Figure 6). Regarding these results, it is rather unlikely that alterations in the nucleoid compactness and the composition of the photosynthetic apparatus are responsible for the reduced growth of barley plants over-accumulating WHIRLY1.

On the other hand, the efficiencies of both photosystems, the maximal electron transport rate (ETR_MAX_), and the quantum yield of photosystem II (Φ(II)) were reduced in plants over-accumulating WHIRLY1. Inversely, the loss of absorbed energy by heat and fluorescence was enhanced in oeW1 plants indicating a malfunctioning of the photosynthetic apparatus. Potentially, changes in the hormone equilibrium could underlie the lower functionality of the photosynthetic apparatus (Muller and Munne-Bosch, 2021, Cackett *et al*., 2022). In this regard, the barley oeW1 plants might be comparable with mutants showing constitutive defense signaling. Arabidopsis mutants with *constitutive expression of pathogenesis-related proteins* (*cpr*) showed a dwarf phenotype (Zhang *et al*., 2003, Heidel *et al*., 2004). To investigate whether the impaired growth is a consequence of deteriorated photosynthesis or energy-consuming defense mechanisms, Mateo *et al*., (2006) investigated the photosynthetic properties of *cpr* mutants in comparison to the WT. Similar to the Arabidopsis *cpr* mutants, barley seedlings overexpressing *WHIRLY1* have a reduced F_V_/F_M,_ a higher ratio of VAZ pool pigments to chlorophylls, and reduced starch content as a consequence of the reduced capacity of the photosynthetic apparatus (Figure 5, 7).

### Overexpression of *WHIRLY1* caused changes in the equilibrium of hormones

Over-accumulation of WHIRLY1 indeed caused a shift in the hormone equilibrium. While the level of the cytokinin isopentenyl riboside (iPR) was reduced, the level of jasmonic acid (JA) is enhanced in oeW1 plants. Cytokinins are well-known for their positive impact on cell division and expansion during leaf development and growth (Brzobohaty *et al*., 1994, Wu *et al*., 2021). Moreover, cytokinins were shown to promote chlorophyll biosynthesis, assembly, and functioning of the photosynthetic complexes (Yaronskaya *et al*., 2006) and to play a role in responses to stress (Albrecht and Argueso, 2017, Cortleven *et al*., 2019). Cytokinins were found to regulate more than 100 genes involved in photosynthesis, including the genes of RubisCO and LHCs (Brenner and Schmulling, 2012) and those encoding sigma factors required for plastid gene transcription by PEP (Danilova *et al*., 2017). Applying cytokinins to wheat leaves increased endogenous cytokinin content and photosynthesis parameters Φ (PSII), F_V_/F_M,_ and ETR, whereas inhibition of cytokinin biosynthesis had opposite effects (Yang *et al*., 2018). The published data suggest that a decreased cytokinin level led to an inactivation of photosystem II reaction centers (Muller and Munne-Bosch, 2021). Hence, the reduced efficiencies of the photosystems, together with the decreased ETR and quantum yield of photosystem II in the oeW1 leaves, are potentially caused by the decrease in the level of iPR. Under HL, cytokinins were reported to promote D1 repair (Cortleven *et al*., 2019). Whereas in the barley plants grown at continuous light of low irradiance, cytokinin levels were low in all genotypes, at high irradiance, the level increased in the WT but not in the oeW1 plants (Figure 8).

Nevertheless, F_V_/F_M_ was higher in the oeW1-14 plants in HL compared to LL. This might indicate that oeW1 plants, compared to WT plants, have a better capacity to respond to HL. A similar finding has been reported for the response of tomato plants overexpressing *WHIRLY1* towards chilling (Zhuang *et al*., 2020b).

In comparison to the cytokinin level, the level of the major auxin IAA was less affected in the barley oeW1 plants. This coincided with similar expression levels of selected genes responding to auxin, i.e. *PIN1* and *TIR1* (Figure S 4). Expression of genes related to auxine biosynthesis such as the YUCCA genes was neither detectable in the WT nor in the oeW1 plants. The level of JA was enhanced in oeW1 plants under standard growth conditions (Figure 8) and during growth in continuous light of low irradiance (Figure S6). It is known that a rise in JA has a negative impact on photosynthesis (Attaran *et al*., 2014, Muller and Munne-Bosch, 2021) and growth (Staswick *et al*., 1992). Recently it has been shown that the treatment of barley leaves with JA affects photosynthesis at the level of the oxygen-evolving complex (Kurowska *et al*., 2020). During growth in continuous light of high irradiance, the level of JA increased in the wild type. As a consequence, the differences among the genotypes measured at low light irradiance disappeared (Figure S6).

In leaves collected under standard growth conditions, the higher expression of defense-related genes such as *PR1* and *HPL*, the latter of which has been proposed as general stress indicator genes (Savchenko *et al*., 2017), indicates that over-accumulation of WHIRLY1 activates defense signaling. Unexpectedly, the level of salicylic acids (SA) stayed below the method’s lowest quantification limit, i.e. 0.1 μM. If the over-accumulation of WHIRLY1 would have induced its synthesis, SA would have increased above a level of 1 μM. The SA-related compounds also did not show changes associated with WHIRLY1 quantities. It may be supposed that SA signaling is not affected by the overexpression of *WHIRLY1* in barley. While in barley, only a limited number of pathogens induced an increase in the level of SA, all tested pathogens induced the expression of PR genes, including *PR1* (Vallelian-Bindschedler *et al*., 1998). Obviously, SA in barley is not always required for defense-related gene expression. In the barley plants grown at continuous high light, also *PR10* expression was enhanced. This gene might be expressed in response to the simultaneous presence of JA and light as reported for rice (Rakwal *et al*., 2001, Zheng *et al*., 2021). A minor contribution of SA to the defense response cannot be excluded considering that expression of *PAL* encoding the key enzyme of salicylic acid biosynthesis is activated in HL both in the WT and much more in the oeW1-50 plants (Figure 10b).

In Arabidopsis, WHIRLY1 was shown to be involved in salicylic acid (SA) signaling independent of NPR1 in the cytosol (Desveaux *et al*., 2004, Vlot *et al*., 2009, An and Mou, 2011, Carella *et al*., 2015). NPR1 is known to translocate from the cytosol to the nucleus upon binding of salicylic acid and thioredoxin-mediated reduction (Mou *et al*., 2003). It has been proposed that WHIRLY1 is translocated from chloroplasts to the nucleus in a similar fashion upon stress-associated redox changes in the photosynthetic apparatus (Foyer *et al*., 2014), whereby the mechanism of translocation remains unknown (Krupinska *et al*., 2022). In the oeW1 plants described in this study, the level of nucleus-located WHIRLY1 is highly upregulated even in the absence of stress. Consequently, in the barley oeW1 plants, defense signaling is constitutively activated, as evident by the expression of *PR1* in fully expanded leaves of seedlings grown under standard growth conditions (Figure 9) and during continuous illumination of low irradiance (Figure 10b). It is obvious that the WHIRLY1-activated defense signaling is mediated by JA rather than by SA. This result is in accordance with reports on JA-dependent defense activation involving PR1 in rice (Yang *et al*., 2013).

By the growth of the oeW1 plants at high irradiance, a dramatic increase in expression of *PR1* (450-fold instead of 70-fold in the wild type) and *PAL* (6-fold instead of only 50% in the WT) was observed (Figure 10b). This indicates that the oeW1 plants are capable of further enhancing defense responses. Considering that the abundance of WHIRLY1 is already high in non-stress conditions, it is unlikely that the higher expression of defense genes is caused by a further increase in WHIRLY1-dependent transcription of these genes. Rather WHIRLY1 abundance may intensify the binding of activating factors to the promoter of *PR1* under certain conditions. Recently, it has been demonstrated that NPR1-mediated *PR1* gene expression requires the formation of an activating complex consisting of histone acetyltransferase (HAC), NPR1, and a TGA transcription factor (Jin *et al*., 2018). Potentially, WHIRLY1 might regulate the accessibility of promoters for defense-associated transcription factors (Krupinska *et al*., 2014a, Krupinska *et al*., 2022a).

Surprisingly, in barley plants overexpressing *WHIRLY1*, the gene encoding isochorismate synthase (ICS) is activated at control conditions. This high expression could be related to a demand for phylloquinone which is essential for electron transfer in photosystem I. A barley *ics* mutant was reported to be deficient in phylloquinone, whereas it was not altered in the basal level of salicylic acid (Qin *et al*., 2019). Salicylic acid biosynthesis may proceed by two possible pathways, the ICS and PAL pathways, which both start from chorismate in chloroplasts (Lefevere *et al*., 2020). In Arabidopsis, only 10% of SA is produced by the PAL pathway, while 90% is produced by the ICS pathway (Garcion *et al*., 2008). By contrast in barley, *ICS* expression during HL exposure is lower than at LL (Figure 10b), while *PAL* expression is enhanced by a factor of 6. This is in accordance with the idea that in barley during stress the PAL pathway of SA biosynthesis is more critical than the ICS pathway. Since, in contrast to *PR1*, the expression levels of *PAL* (Figure 10) and of the defense gene *THIO1* (Figure S4) were not elevated by overexpression of WHIRLY1 under normal growth conditions, it is unlikely that these genes are directly regulated by WHIRLY1. In contrast, *PR1* and *HPL* were activated in the oeW1 plants both under normal growth conditions and at HL and, therefore might be directly activated by WHIRLY1.

### The role of chloroplast-nucleus located WHIRLY1 in the growth-defense tradeoff

The reduced growth of oeW1 plants and the enhanced resistance towards powdery mildew indicate that overexpression of *WHIRLY1* shifts the balance between growth and resistance to the latter. In recent years hormone crosstalk has emerged as a major player in regulating the growth-defense tradeoff (Huot *et al*., 2014). Although the antagonistic crosstalk between SA and the growth hormone auxin mostly has been reported to determine the tradeoff between growth and defense (Huot *et al*., 2014), overexpression of *WHIRLY1* in barley had more impact on the levels of cytokinins and JA than on those of auxin and SA, suggesting that in this species the tradeoff is regulated by cytokinin and JA However, most studies on hormonal interactions during growth and defense have been performed with Arabidopsis. It is likely, that hormonal interactions in monocot plants are different, as has been reported for rice (De Vleesschauwer *et al*., 2013). It has been postulated that during immune responses in rice, NPR1-dependent SA-signaling is activated by JA binding to the COI1 receptor without a change in the level of SA (Yang *et al*., 2013). This model is in accordance with earlier reports on barley infection by powdery mildew, in which sensitivity to powdery mildew was found to be not accompanied by accumulation of SA (Vallelian-Bindschedler *et al*., 1998, Hückelhoven *et al*., 1999).

The high accumulation of WHIRLY1 in chloroplasts of the barley oeW1 plants had neither consequences for nucleoid organization nor plastid gene expression. Hence the reduced growth was likely not caused by changes in the plastid gene expression machinery but rather by the enhanced level of nucleus-located WHIRLY1, inducing changes in gene expression that eventually lead to a rewiring of hormonal homeostasis. The identical molecular weights of chloroplast-located WHIRLY1 and nucleus-located WHIRLY1 clearly indicate that both pools of WHIRLY1 had been processed to the mature form inside chloroplasts. Hence WHIRLY1 over-accumulating in the nucleus was transferred from chloroplasts to the nucleus as demonstrated before with transplastomic tobacco plants synthesizing WHIRLY1 inside chloroplasts (Isemer *et al*., 2012b). In another previous study, it had been shown that Arabidopsis plants accumulating WHIRLY1 outside the chloroplasts behave like a WHIRLY1-deficient mutant (Isemer *et al*., 2012a). Hence, the nuclear activities of WHIRLY1 require its preceding presence in chloroplasts. Whether WHIRLY1 undergoes a modification inside chloroplasts and how its transfer to the nucleus is mediated remains to be determined. Taken together, the findings of this study suggest that the WHIRLY1-mediated adjustment of hormonal homeostasis is controlled by chloroplasts which are crucial sensors of environmental information (Pfalz *et al*., 2012, Zhang *et al*., 2020).

According to the elevated *PR1* expression in the absence of stress, barley oeW1 plants showed a constitutive defense response. To avoid a negative impact on growth, the expression of resistance genes might be restricted to the time of stress perception and the subsequent defense response (Karasov *et al*., 2017). Sequestering of WHIRLY1 in chloroplasts is a means to avoid its nuclear activity under non-stress conditions and to allow a fast response to stress only under conditions that induce the transfer of WHIRLY1 from chloroplasts to the nucleus (Krause and Krupinska, 2009). However, the 10 to 50 times higher level of WHIRLY1 in the oeW1 plants obviously exceeded the capacity for WHIRLY1 sequestration by chloroplasts. It remains to be tested whether a moderate increase in WHIRLY1 accumulation in the chloroplast is possible without transfer to the nucleus in non-stress conditions, thereby avoiding the constitutive expression of *PR1* in the nucleus.

## EXPERIMENTAL PROCEDURES

### Plant material and growth conditions

Transgenic barley plants overexpressing *HvWHIRLY* under the control of the maize *UBIQUITIN 1* promoter were generated by the transformation of barley immature embryos by *Agrobacterium tumefaciens* as described (Hensel *et al*., 2008). The pENTR/TOPO Gateway vector (Invitrogen, Karlsruhe, Germany) was used for the transfer to the pIPKb007 binary vector using Gateway™ LR as described (Himmelbach *et al*., 2007). Plantlets resistant to hygromycin were transferred into soil and cultivated in a greenhouse. Additionally, PCR analyses with primers (Krupinska et al. 2014, Supplementary Table 1) for the hygromycin resistance cassette were performed to verify the transgene integration. As control plants, the barley cultivar “Golden “Promise’ and for powdery mildew assays, the *HvWHIRLY1* knockdown plants (RNAiW1-7) (Krupinska *et al*., 2014b) were used.

Barley grains were sown on soil (Einheitserde ED73, Tantau, Ütersen, Germany) and transferred for three days in a dark and cold chamber (6°C) to synchronize germination. Thereafter, the grains were transferred to a climate chamber where the seedlings were grown either in a standard daily light/dark cycle (16:8) as described (Krupinska *et al*., 2019) or in continuous light of different irradiances (100 or 350 μmol photons m^-2^ s^-1^) as also described previously (Swida-Barteczka *et al*., 2018).

### Quantum yields of the photosystems and electron transport rate

The maximum quantum yield of photosystem II, F_V_/F_M_, and the maximum P700 (P_M_) signal were measured in parallel by Dual-PAM-100 (Walz GmbH, Effeltrich, Germany). The leaves were kept for about 10-15 minutes under low light (20-40 μmol m^-2^ s^-1^) before starting the measurement. The measurement was done at 13 different light levels, starting from zero and gradually increasing during six minutes to 1600 μmol m^-2^ s^-1^. In between of these light levels, there was a step with 60 μmol m^-2^ s^-1^ which is similar to the growth light in the climate chamber. The quantum yields of photosystem II as well as of non-radiative and radiative dissipation were calculated as follows (Klughammer and Schreiber, 2008): Ф(II) = (*F*_*M*_′−*F*)/*F*_*M*_′, Ф(NPQ) = F/*F*_*M*_′- F/F_M,_ Ф(NO) = F/F_M._

### Determination of pigments by high-performance liquid chromatography

For the analysis of pigments, one cm long leaf segments excised from the area between 1.5 and 3 cm below the leaf tip were immediately frozen in liquid nitrogen and kept at −80°C. Pigments were extracted and HPLC analysis was performed as described (Saeid-Nia *et al*., 2022). To calibrate the detector (Nichelmann *et al*., 2016), pure carotenoid extracts (except antheraxanthin) were prepared through thin-layer chromatography (modified after Lichtenthaler and Pfister 1978). Afterwards, the concentrations of the pure pigment solutions were determined by spectrophotometry using the extinction coefficients provided by Davies (1976).

### Immunoblot analyses

Total proteins were extracted from ground leaf material and subjected to SDS-PAGE, as reported (Krupinska *et al*., 2014, 2019). Proteins were transferred onto the nitrocellulose membrane by semi-dry electroblotting. Antibodies against PsaA (AS06172), PsbA/D1 (AS01016), LHCA1 (AS01005), and LHCB1 (AS01004) were purchased from Agrisera. The antibody directed towards HvWHIRLY1 was prepared against a synthetic peptide and can be purchased from Agrisera (AS163953). Immunoreactive complexes were visualized using a peroxidase-couples secondary antiserum with chemiluminescence detection kits (ECL Select, Amersham, USA; Lumigen, Southfield, MI, USA). The ChemiDoc MP Imaging Systems and the Image Lab 6.1 software (Bio-Rad Laboratories, Munich, Germany) were used for the quantification of signal intensities.

### Determination of hormones

Leaf samples of ca. 30 mg (fresh weight) were weighed into 2 ml safe lock tubes (Eppendorf AG, Germany) and kept at -80°C until analysis. Empty tubes were used as blanks. Before extraction, two 3 mm ceria-stabilized zirconium oxide beads were placed into each tube. The samples were extracted and purified as described by Šimura *et al*. (2018) with minor modifications (Simura *et al*., 2018). The absolute quantification of all targeted phytohormones, excluding salicylates, was performed as described (Eggert and von Wiren, 2017).

The analysis of salicylates was performed using UHPLC-HESI-HRMS (Vanquish UPLC) coupled to QExactive Plus Mass Spectrometer (San Jose, CA, USA). The MS was equipped with a HESI source operating in negative ion mode. Salicylates baseline separation was achieved on a reversed-phase Acquity UPLC® HSS T3 column (10 Å, 2.1 × 100mm, 1.8μm, Waters) using a gradient elution of A (Water, 0.1% FA) and B (ACN, 0.1% FA) as follows: 0– 5min, 5% B; 5–10min, 5% to 80% B. Additional five minutes were added for column washing and equilibration (total run time, 15min). The column temperature was set at 45°C and the flow rate at 0.5 ml·min-1. The injection volume was 5μl. Source values were set as follows: Spray voltage 2.5kV; capillary temperature 255°C; S-lens RF level 40; Aux gas heater temp 320°C; Sheath gas flow rate 47; Aux gas flow rate 11. For spectra acquisition, a Full MS/dd-MS^2^ experiment was performed. Resolution in Full Scan was set as 70000. For MS/MS experiments, resolution 17,500 and NCE 40V were used. The identification of compounds found in extracts was based on a comparison of their retention times, MS2 spectrum and exact mass with standards.

### RNA isolation and real-time PCR analysis

Total RNA was isolated from primary foliage leaves of seedlings using the peqGOLD-TriFast reagent (Peqlab Biotechnology, Erlangen, Germany) according to the manufacturer’s protocol. cDNA biosynthesis and real-time PCR were performed as described previously (Krupinska *et al*., 2019). Data were normalized to the level of the ADP-ribosylation factor 1 mRNA (Rapacz *et al*., 2012), to cytosolic GAPDH or to mRNA of the barley histone acetyltransferase (HORVU.MOREX.r2.1HG0027750), which has the alternative name GENERAL CONTROL NONREPRESSIBLE 5 (GCN5).

### Transmission electron microscopy

Leaf segments from primary foliage leaves (2×2mm) at a position of 2 cm below the leaf tip were fixed and processed as described (Krupinska *et al*., 2014b).

### Staining and localization of nucleoids

Leaf cross-sections were produced by hand or by a hand microtome from the primary foliage leaf of plants grown for 7 days. Sections were fixed by 4% (w/v) paraformaldehyde in phosphate-buffered saline (PBS) overnight at 4°C. After washing with PBS containing 0.12 %(w/v) Glycin, the sections were stained with SYBR®Green (1:5000, S7563 InvitrogenTM) for 45 min in darkness at room temperature. After washing with 1x PBS for 15 min, the sections were transferred onto a slide, capped with PBS/glycerol (v/v: 1:1), and a coverslip. Imaging was done at Leica SP5 confocal microscope system with an HCX PL APO CS 63.0 × 1.2 W objective. Excitation was done by an argon laser line 488 (5% power). Emission was detected between 510-570 nm (HV750) and 690-760 nm (HV480). A minimum of five images out of different regions of the specimen were taken from each sample. Image analysis, coloring, and composition were done by ImageJ 1.53q.

### Infection with powdery mildew

Five plants were grown in 12 cm pots in compost soil. In an inoculation device, transgenic lines with two pots each were arranged, with three pots containing wild type. While rotating in the inoculation tower, the fourteen-day-old seedlings were inoculated with *Blumeria graminis* spores (isolate CH4.8) until a spore density of approx. 10 spores per mm^2^ have been reached. The disease scored 7 d after inoculation, as described (Schweizer *et al*., 1995).

## ACKNOWLEDGEMENTS

We thank Sabine Sommerfeld (IPK Gatersleben) and Susanne Braun (Institute of Botany, CAU, Kiel) for their excellent technical assistance. We are grateful to the German Research Foundation for financial support (KR1350-19-1).

## SUPPORTING INFORMATION

Additional Supporting Information may be found in the online version of this article:

**Figure S1**. (a) The abundance of WHIRLY1 in primary leaves of the lines oeW1-2 and oeW1-14. (b) Immunoblots showing the distribution of WHIRLY1 between chloroplasts and nucleus.

**Figure S2**. Relative levels of mRNAs of *HvWHIRLY1* and selected plastid genes (*rpoB, clpP, psaA, psbA, psbE*) were determined by RT-PCR.

**Figure S3**. Nucleoid morphology in different parts of primary foliage leaves from wild type, oeW1-50 and oeW1-10 plants.

**Figure S4**. Expression of *PIN1, TIR1, THIO1*and *GR1* in primary foliage leaves from oeW1-2 and oeW1-14 grown in a daily light/dark cycle.

**Figure S5**. Expression of defense genes in primary foliage leaves from oeW1-2 and oeW1-14 grown in continuous light of low (LL) or high irradiance (HL).

**Figure S6**. Hormone levels supplementing Figure 8. (a) Levels of SAG and DHBA in primary foliage leaves from oeW1-2 and oeW1-14 grown in a daily light/dark cycle, (b) levels of iRP, IAA and JA in primary foliage leaves from oeW1-2 and oeW1-14 grown in continuous light of low (LL) or high irradiance (HL).

