## Supplemental Information for "The balance between growth and resistance is shifted to the latter by over-accumulation of chloroplast-nucleus located WHIRLY1 in barley"

*[#]current address: Institute of Plant Pathology, University of Bonn, Germany*

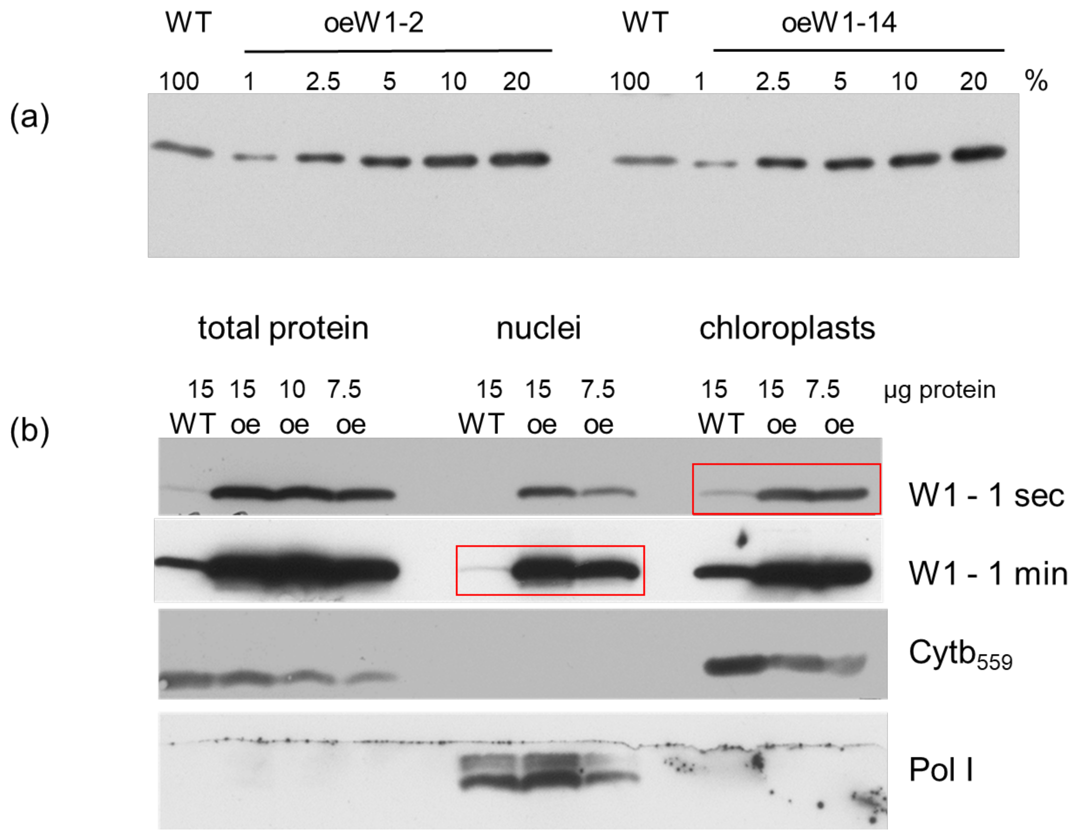


**Figure S1.** Abundance of WHIRLY1 in primary leaves of the lines oeW1-2 and oeW1-15 in comparison to the wild type. (a) To elucidate the relative amount of WHIRLY1 in oeW1-2 and oeW1-14 leaves, different protein amounts were loaded and compared to the wild-type sample (100%). (b) Relative abundance of WHIRLY1 in chloroplasts and nuclei prepared from primary foliage leaves of oeW1-2. In case of the oeW1-2 samples different amounts of protein were loaded. After incubation with the WHIRLY1 antibody the blot was exposed to film for different times (1 sec, 1 min). The frames allow a visual comparison between relative abundance of WHIRLY in chloroplasts and nucleus. The purity of fractions has been tested by immunodetection of chloroplast located cytochrome $b_{559}$ and nucleus located DNA polymerase I (Pol I).

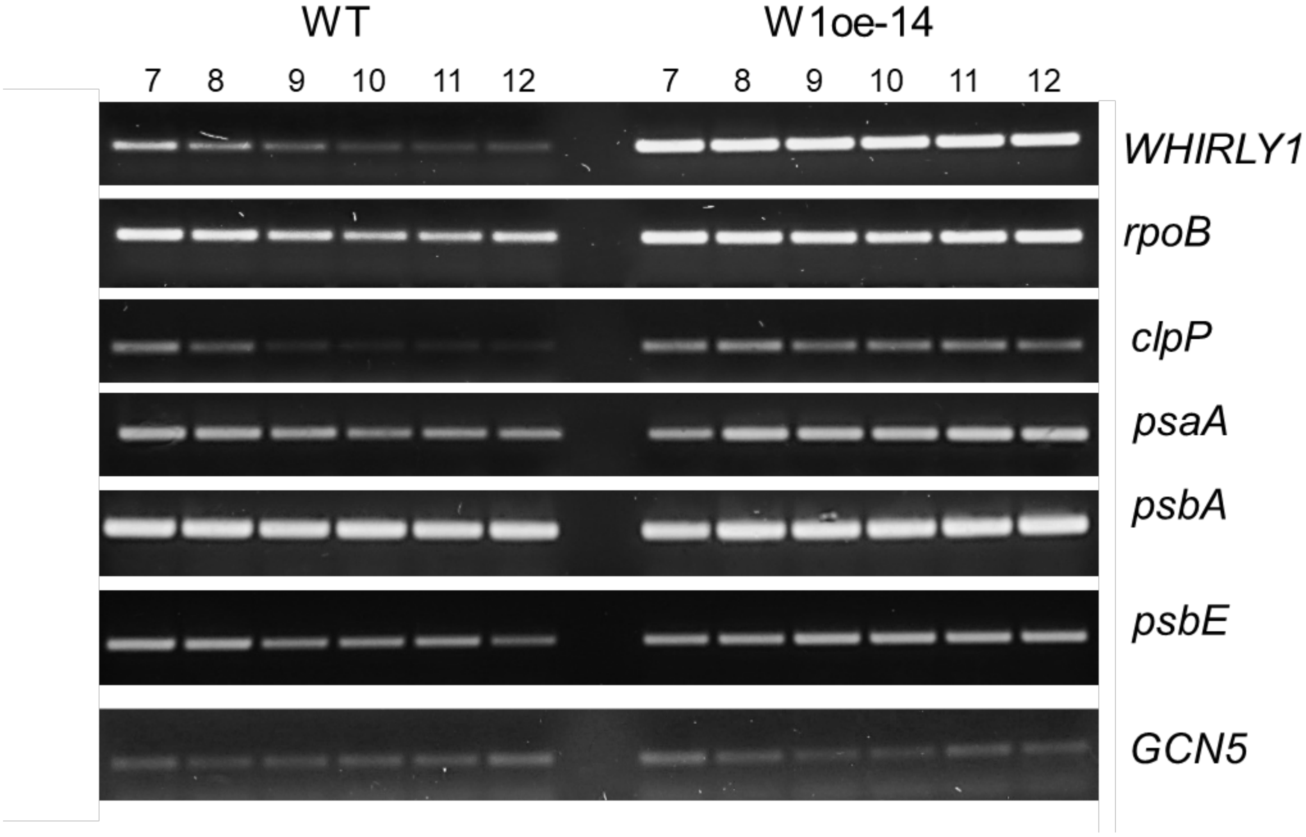


**Figure S2.** Relative levels of mRNAs of *HvWHIRLY1*and selected plastid genes (*rpoB, clpP, psaA*, *psbA, psbE*) determined by RT-PCR. The relative mRNA levels were compared between primary foliage leaves of the wild type and oeW1-14 at 7-12 das. As a constitutively expressed standard gene *GCN5* was used.

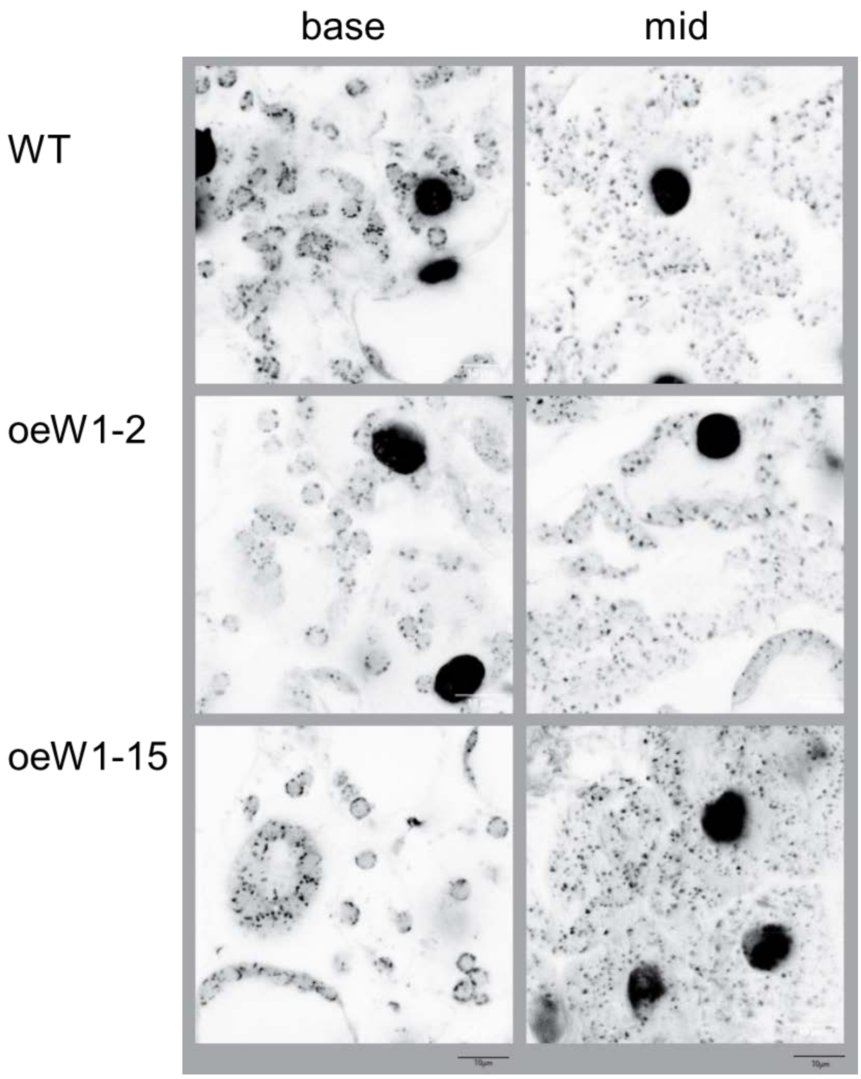


**Figure S3.** Developmental changes in the morphology of plastid nucleoids in wild type, oeW1-2 and oeW1-15 seedlings. Sections from the base, the mid part and the tip of primary foliage leaves were stained with SYBR Green. Excitation was done by an argon laser line 488 (5% power). Emission was detected between 510-570 nm (HV750) and 690-760 nm (HV480).

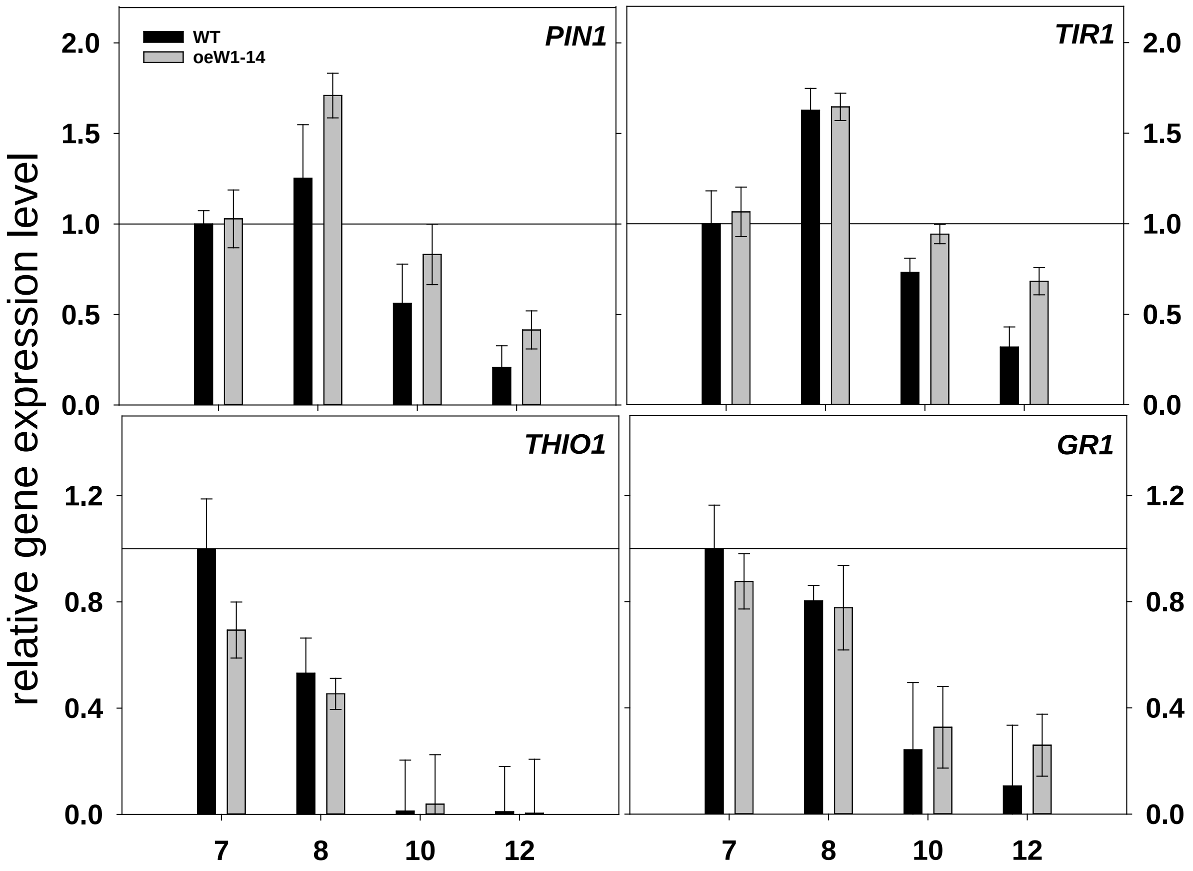


**Figure S4.** Expression of *PIN1, TIR1, THIO1* and *GR1* measured by qRT-PCR using *GCN5* (see Material and Methods) as standard. RNA was extracted from primary foliage leaves of wild-type and oeW1-14 seedlings grown for different times (7, 8, 10 and 12 das) in a daily light/dark cycle. Columns are means +/- standard deviations of n=3 (10 pooled leaves).

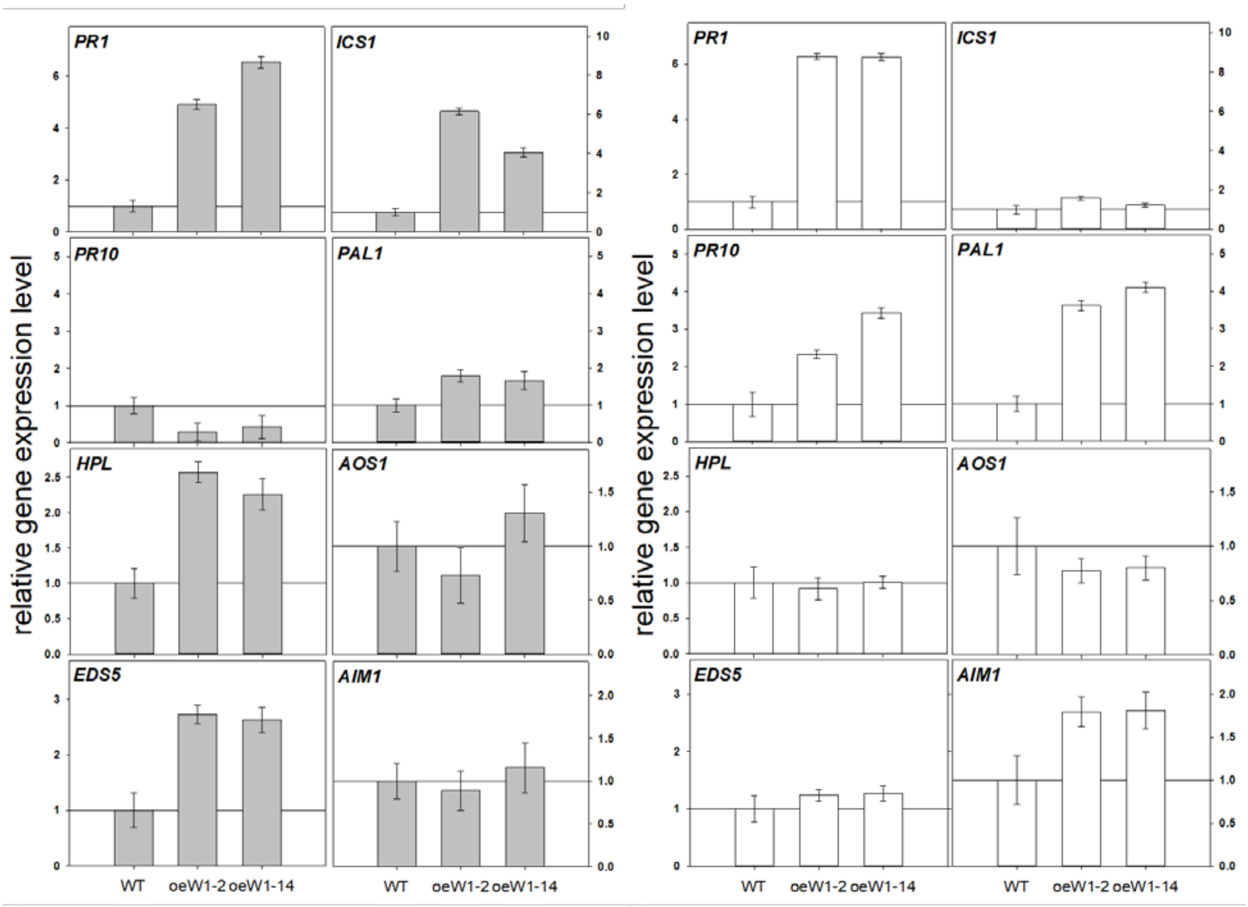


**Figure S5.** Gene expression in primary foliage leaves of wild type and oeW1 seedlings of lines with a 50-fold higher abundance of HvWHIRLY1 (oeW1-2, oeW1-14) during growth in continuous light of low (left) and high irradiance (right). Genes: *PR1, ICS, PR10, PAL1, HPL, AOS1, EDS5, AIM1* expression of defense-related genes putatively associated with high light stress and SA: *PR1, PR10, ICS, PAL, HPL, PAL1, AOS.*

**(a)**

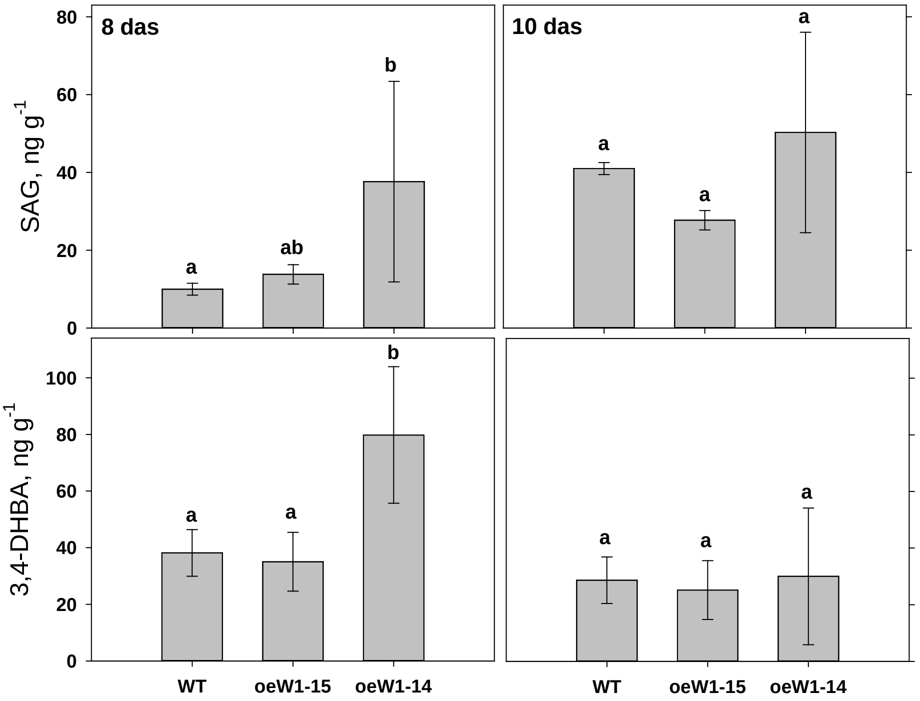


**(b)**

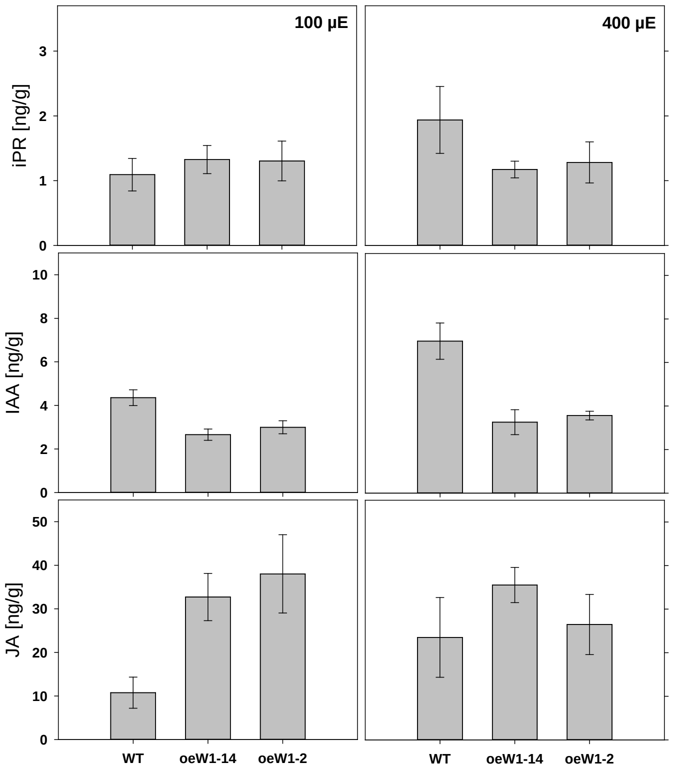


**Figure S6.** Hormone levels in primary foliage leaves of the wild type in comparison to oeW1 seedlings. (a) Seedlings of WT, oeW1-15 and oeW1-14 lines were grown in a daily light/dark cycle and leaves were collected at different days after sowing (8 and 10). (b) Seedlings of two oeW1 lines (oeW1-2, oeW1-14) were grown in continuous light of low (100 µE) or high irradiance (400 µE). SAG=salicylic acid glucoside, DHBA=dihydroxy benzoic acid, iPR=isopentenyl riboside, IAA=indole acetic acid, JA=jasmonic acid. Columns are means +/- standard deviations of n=5 leaves.
